# Old foe, new host: epidemiology, genetic diversity and pathogenic characterization of maize streak virus in rice fields from Burkina Faso

**DOI:** 10.1101/2023.10.02.560455

**Authors:** Noun Magdy Ibrahim Fouad, Mariam Barro, Martine Bangratz, Drissa Sérémé, Denis Filloux, Emmanuel Fernandez, Charlotte Julian, Nignan Saïbou, Abalo Itolou Kassankogno, Abdoul Kader Guigma, Philippe Roumagnac, Issa Wonni, Charlotte Tollenaere, Nils Poulicard

## Abstract

Rice is of critical significance regarding food security worldwide including in Africa. Only two viruses impacting rice production in Africa have been deeply investigated for decades: the rice yellow mottle virus (*Solemoviridae*) and the rice stripe necrosis virus (*Benyviridae*). Using viral metagenomics, we aimed at exploring the diversity of viruses circulating in Burkina Faso rice fields. We performed an epidemiological survey in this country between 2016 and 2019 involving 57 small farmer’s rice fields under two production systems (rainfed lowlands and irrigated areas). More than 2700 rice samples were collected without *a priori* (not based on symptom observation) following a regular scheme. In addition, wild and cultivated (maize and sugarcane) *Poaceae* growing nearby rice fields were also collected. Unexpectedly, metagenomics detected maize streak virus (MSV, *Geminiviridae*) in analyzed rice samples. Further molecular analyses using RCA-PCR showed that MSV is widely distributed and highly prevalent in both rainfed lowlands and irrigated rice areas. MSV-A and MSV-G strains were identified. MSV-G, exclusively identified so far in wild grasses, was the most prevalent strain while MSV-A, known to cause severe symptoms in maize, was sporadically identified. No genetic differentiation was detected between MSV isolates either infecting wild or cultivated plant species. Using infectious clones in experimental conditions, we confirmed the pathogenicity of both MSV strains on rice. Thus, in addition to contribute to the epidemiological surveillance of rice production in Africa, our results illuminate new epidemiological and pathogenic aspects of one of the most studied plant viruses with significant economic consequences in Africa.

**Funding:** French National Research Agency «Investissements d’avenir» program (ANR-10-LABX-001-01), Agropolis Fondation (ANR-16-IDEX-006), French National Research Agency “young researchers” program (ANR-20-CE35-0008-01), CGIAR Research Program on Rice Agri-food Systems (RICE), Cooperation and cultural action department of the French Embassy (SCAC) in Burkina Faso.

## INTRODUCTION

Emergent crop diseases, a high proportion of which are caused by viruses, are a significant burden on food security and economic stability of societies, especially in developing countries (Anderson et al. 2004; Jones 2021). While the intensification of agriculture has become one of the major priorities to provide food for people, the intensification programs are threatened by climate and global changes (environmental, demographic and socio-economic) which could subsequently favor the emergence of plant pathogens and increase their impact on food production (Anderson et al. 2004; Baker et al. 2022).

Rice is the most important human staple food crop in the world, directly feeding nearly half of world’s population every day. In Africa, rice cultivation has historically involved two species: the African rice *Oryza glaberrima* that was domesticated in West Africa ca. 3000 years ago and the Asian rice *O. sativa* that was introduced repeatedly since the 15th centuries (Portères 1970). From the second half of the 20^th^ century and more intensively during the last decades, cultivated rice areas have drastically increased in Africa and this crop has become important as a strategic commodity for food security, in particular for facing the effects of recent acute demographic and societal changes (FAOSTAT 2022, https://data.un.org/Data.aspx?d=FAO&f=itemCode%3A27; Demont 2013; Soullier et al. 2020). Such agricultural changes, including the preference and intensification of the Asian rice cultivation (Cubry et al. 2018), could render the rice cultivation more exposed to pathogen emergences and epidemics, particularly to viruses (Anderson et al. 2004). Our capacity to ensure the sustainability of production systems against plant diseases strongly depends on our ability to explore the vast diversity of microorganisms in the environment, to understand how they interplay with the stability and productivity of ecosystems, and to identify pathogens from their early stage of emergence thought epidemiological surveillance.

Among the 19 viruses that have been so far reported to infect rice worldwide (Wang et al. 2022), five viruses have been occasionally identified in Africa (Abo and Sy 1997). Only two of these five viruses detected in Africa have been genetically characterized and deeply investigated, the rice stripe necrosis virus (RSNV, *Benyvirus*, *Benyviridae*; Bagayoko et al. 2021) and the rice yellow mottle virus (RYMV, *Sobemovirus*, *Solemoviridae*; Hébrard et al. 2021). The three remaining viruses of rice reported in Africa that were poorly characterized so far are the maize streak virus (MSV strain A, *Mastrevirus*, *Geminiviridae*), the African cereal streak virus (ACSV, genus and family not determined) and the rice crinkle disease (aetioloogy, genus and family not determined; Abo and Sy, 1997).

The recent methodological innovations in high-throughput sequencing (HTS) and metagenomics give us the opportunity to explore the vast genetic and functional diversity of viruses in wild and cultivated environments and to identify putative emergent viruses (Bernardo et al. 2018; Edgar et al. 2022; Greninger 2018; Lefeuvre et al. 2019). However, despite this continuous detection of partial or complete virus metagenome-assembled genomes (MAGs) from wild or anthropized environments, few of these MAGs have subsequently been subjected to more in-depth analyses, in most cases using Sanger sequencing. In addition, this sequencing validation step has usually not been supplemented with epidemiological or pathogenetic information on the related viruses (Koonin and Dolja 2018). Nevertheless, among the few novel viruses initially discovered using metagenomics-based approaches and further partially or fully biologically characterized, geminiviruses were among the most represented (see Table 4 in Moubset et al. 2022).

In this study, we intended to meet these expectations focusing on viruses circulating in rice fields in Burkina Faso. For that, an epidemiological survey focusing on two rice production systems (irrigated and rainfed lowland) was conducted in this West African country between 2016 and 2019 (Figure 1). In parallel, agricultural practices and the use or turnover of rice varieties were followed as well as the diversity of microbiome associated to rice roots and (known) rice pathogens (viral, bacterial and fungal) circulating in these fields (Barro et al. 2022, 2021a, b; Billard et al. 2023; Kaboré et al. 2022; Tollenaere et al. 2017). Based on these samples and metadata, we performed viral metagenomics analyses on rice and wild grass leaf samples collected in 2016-2017 (project MP1 in Moubset et al. 2022). Among the thousands of contigs assigned to viral genomes (including RYMV), these analyses revealed unexpected virus species infecting rice as maize streak virus (MSV, *Mastrevirus*, *Geminiviridae*).

**Figure 1:**
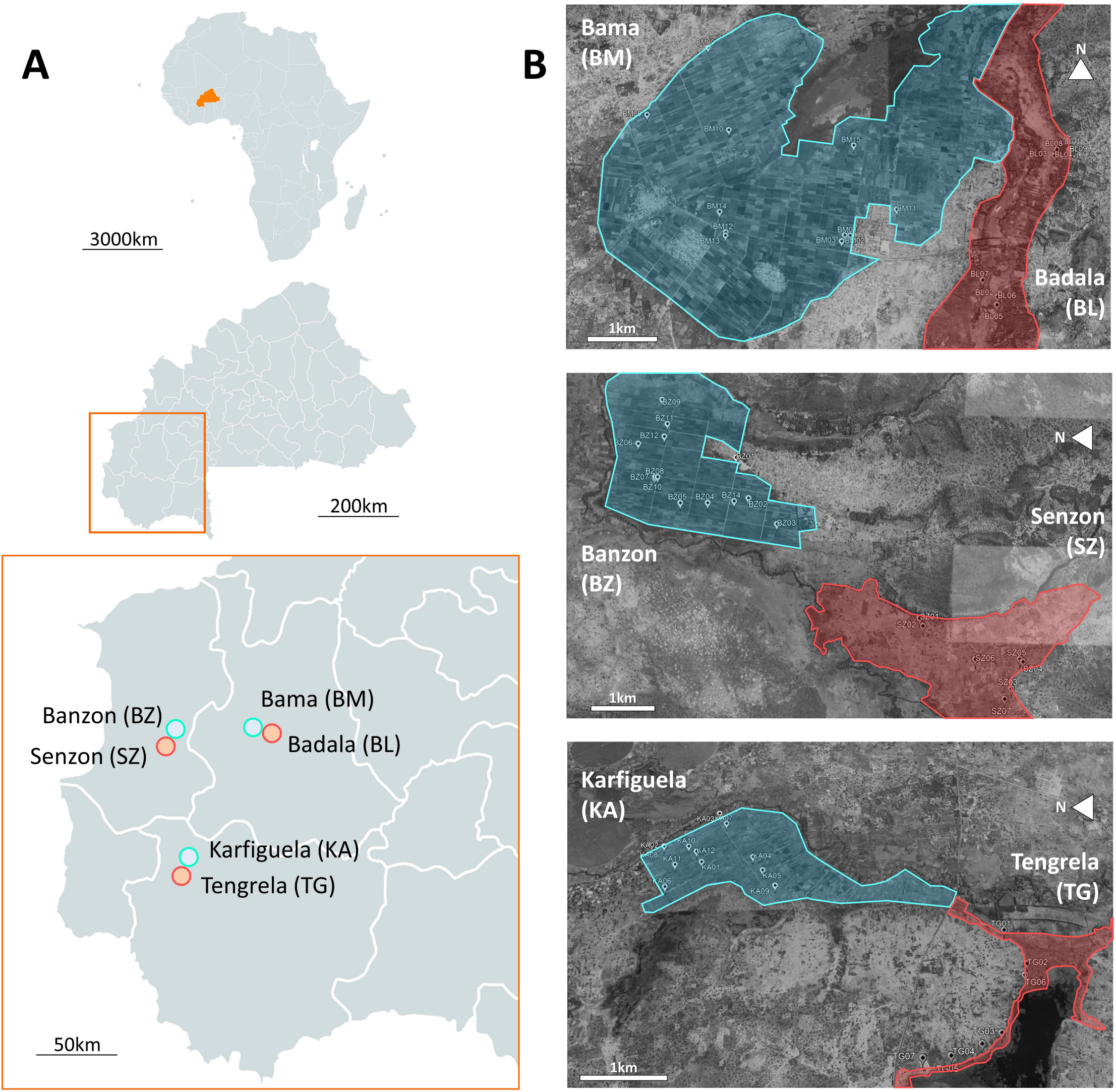
Study area for the spatiotemporal survey of rice fields in Burkina Faso. **A**. Burkina Faso location in Africa, with a focus on western Burkina Faso where are located the six sites: Bama (BM in the text), Badala (BL), Banzon (BZ), Senzon (SZ), Karfiguela (KA) and Tengrela (TG). These sites are indicated with colored dots referring their rice production systems: irrigated (IR, blue) and rainfed lowland (RL, red). **B**. Distribution area of irrigated (blue) and rainfed lowland (red) rice production systems in the six sites and location of the rice fields surveyed during this study.

Geminiviruses are responsible for a large number of emerging crop diseases in the World, with considerable impact on the yields of several cash and staple crops (maize, cassava, tomato, cotton, beans and grain legumes) and constitute a major threat to the food security of tropical and sub-tropical developing countries (Rybicki 2015; Rybicki and Pietersen 1999). Specifically, maize streak virus (MSV), one of the most devastating viruses on maize in Africa (Martin and Shepherd 2009; Savary et al. 2019) was first described in South Africa during the 1870’s and later in 1896 during a serious outbreak (Fuller 1901; Shepherd et al. 2010). MSV virion consists in twinned and quasi-icosahedral (geminate) capsid. Its monopartite circular single-stranded DNA (ssDNA) genome of ca. 2.7kb, replicating by rolling circle encodes only four proteins (Fiallo-Olivé et al. 2021; but see Gong et al. 2021). Bidirectional transcription from a long intergenic region (LIR), specifically from the highly conserved origin of replication at the top of a stem-loop structure of ca. 50 nucleotides (Heyraud et al. 1993), results in the virion sense expression of a movement protein (MP) and a coat protein (CP) and the complementary sense expression of the replication-associated proteins, Rep and RepA (Fiallo-Olivé et al. 2021). MSV is transmitted naturally by 18 leafhopper species (Fiallo-Olivé et al. 2021), some of which have been observed in the study area (personal communication) and reported in rice fields of this region (Tra Bi et al. 2020). Alternatively, MSV is also transmitted experimentally by agroinoculation of infectious clones (Boulton et al. 1989; Grimsley et al. 1987). In addition to maize, MSV is characterized by a large monocotyledonous host range including wild and cultivated*Poaceae* species, such as sugarcane and sorghum (Kraberger et al. 2017). Restricted to Africa and surrounding islands, 11 strains (from A to K) have been identified. MSV-A is the only strain known to cause severe symptoms on cultivated plants, whereas MSV-B, -C and -E, mainly detected in infected wild grasses, can also produce mild infections in MSV-susceptible maize genotype (Claverie et al. 2019; Oyeniran et al. 2021; Shepherd et al. 2010). Previous studies showed that, as with other geminiviruses, recombination played a decisive role in the evolution of MSV. Particularly, a recombination event that occurred in the mid-19^th^ century between ancestral MSV-B and MSV-G/-F strains probably led to the emergence, the efficient dispersion across Africa and the progressive adaptation of MSV-A strain to maize that became a major pathogen to this crop (Harkins et al. 2009; Monjane et al. 2020, 2011; Varsani et al. 2008).

While MSV-A strain has recurrently been reported in maize from Burkina Faso (Kraberger et al. 2017), it was only identified once in rice fields in this country (Konaté and Traoré 1992). Noteworthy, MSV-A strain was also experimentally transmitted to rice by viruliferous leafhoppers (Damsteegt 1983; Konaté and Traoré 1992). However, information on geographical distribution, prevalence, genetic diversity and aggressiveness of this virus in rice fields from Burkina Faso has never been reported to our knowledge. Thus, based on viral metagenomics, epidemiological and experimental approaches, we intended in this study to evaluate the epidemiological and pathogenic status of MSV in rice fields from this country. In addition to contribute to the epidemiological surveillance of rice production in Africa, our results participate to illuminate new epidemiological and pathogenic aspects of one of the most studied plant viruses with significant economic consequences in Africa.

## MATERIAL AND METHODS

### Study area and samplings

The study area is located in western Burkina Faso, in a 100 x 100-km region in the Sudanian bioclimatic area (Figure 1A; map based on MapChart website, https://mapchart.net/). Six sites located within three geographical zones (Bama-BM/Badala-BL, Banzon-BZ/Senzon-SZ and Karfiguela-KA/Tengrela-TG) have been surveyed between 2016 and 2019. Each geographical zone comprised one irrigated (IR) site and a neighboring rainfed lowland (RL) site (Figure 1B; field locations and delimitations based on GoogleEarth maps).

A regular and longitudinal rice leaves sampling in these six sites involving 57 rice fields have already been performed as previously described in Barro et al. 2021a). Briefly, observations and samplings were assessed at the maximum tillering/heading initiation stages, from September to December each year. Each studied field was approximately a square of 25 meters on each side, with a regular sampling of 16 plants per field over a grid (Figure Supp1, https://doi.org/10.23708/8FDWIE). This sampling approach did not consider the potential disease symptoms (regular sampling without *a priori*), but the rice leaves were inspected when sampled and disease symptoms were recorded when observed.

In addition to these rice samples, the diversity and the percentage of surface covering of wild grass species growing in the rice field borders were estimated for 6 fields in 2017 (irrigated production system: BM02, BZ11, KA01; rainfed lowland production system: BL02, SZ07, TG01; Figure Supp1). One plant of the five most frequent plant species was randomly collected without *a priori* (*i.e.* independently to observation of disease symptoms) for further analyses (see below). Finally, specific samplings of wild (*Poaceae* species) or cultivated (rice, maize and sugarcane) plants presenting symptoms putatively related to virus infection (leaf deformations, stripes, …) was performed within or nearby the rice fields involved in the longitudinal survey (Figure Supp1).

Thus, the total number of plant samples analyzed during this study is above 2800, with ca. 2750 rice plant samples (from 43, 42, 49 and 40 rice fields in 2016, 2017, 2018 and 2019, respectively) and 30 wild grasses collected without *a priori*, and 4, 8, 7 and 15 symptomatic samples of rice, maize, sugarcane and wild *Poaceae*, respectively. These samples were therefore named according to the date, the site, the field and the host plant of collection (*i.e.* 17TG01 and 17TG01w correspond respectively to rice and wild plant samples collected in 2017 in/close to the rice field “01” of Tengrela).

### Viral metagenomics

Detection and identification of both DNA and RNA viruses were performed on rice and wild *Poaceae* samples using virion-associated nucleic acid (VANA) metagenomics-based approach (Moubset et al. 2022). Specifically, for each rice field surveyed in 2016 and 2017, a pooled sample of 1g of the 16 sampled rice plants was grinded and prepared for analysis (*i.e.* 85 pooled rice samples were obtained to represent 85 rice fields: 43 fields in 2016 and 42 in 2017). Similarly, the 5 leaf samples of the 5 most frequent wild plant species collected without *a priori* in each border of rice field were grinded and pooled (*i.e.* 6 pooled grass samples were obtained to represent 6 rice field borders surveyed in 2017).

Each pooled sample of rice or wild grasses were processed using the VANA approach as described by François et al., 2018). Briefly, we isolated viral particles by filtration and ultracentrifugation. The nucleic acids not protected in virus-like particles were further degraded by DNase and RNase and then the total RNAs and DNAs were extracted using a NucleoSpin kit (Macherey Nagel, Bethlehem, PA, USA). Reverse transcription was performed by SuperScript III reverse transcriptase (Invitrogen), cDNAs were purified by a QIAquick PCR Purification Kit (Qiagen, Hilden, Germany) and complementary strands synthesised by Klenow DNA polymerase I. Double-stranded DNA was amplified by random PCR amplification. Samples were barcoded during reverse transcription and PCR steps using homemade 26 nucleotide (nt) Dodeca Linkers and PCR multiplex identifier primers. PCR products were purified using NucleoSpin gel and PCR clean-up (Macherey Nagel, Bethlehem, PA, USA). Finally, libraries were prepared from purified amplicons and sequenced on an Illumina HiSeq to generate 2x150nt paired-end reads (Genewiz, South Plainfield, NJ, USA).

### Nucleic acid extractions, rolling circle amplification and MSV detection by specific PCR

Total DNA extraction of each sample (100mg of plant material for pooled leaf samples, 20mg for individual leaf sample) was performed according to the CTAB protocol and the concentration and quality of extracted DNA were assessed with NanoDrop Microvolume Spectrophotometers (ThermoFisher Scientific, Waltham, MA, USA).

The circular DNA genomes of MSV were first amplified by rolling circle amplification (RCA) according to the manufacturer’s protocol (TempliPhi^TM^ kit, GE Healthcare, Munich, Germany). Then, the presence of MSV-G and MSV-A at field or plant level was detected by PCR using 1µl of RCA products, GoTaq Flexi (PROMEGA, Madison, WI, USA) according to the manufacturer’s protocol and primers targeting MSV-G (MSV-F559bp: 5’-GGAGCATGTAAGCTTCGGGA-3’, positions 1875-1889; MSV-R559bp: 5’-GAGCTCGTTGGTCACTGGAA-3’, positions 2415-2434, Tm=57°C, amplification: 559bp) and MSV-A (MSVg-2F: 5’-TCAGCCATGTCCACGTCCAAG-3’, positions 478-498; MSVa-1R: 5’-TCACCACGAAGCGATGACACA-3’, positions 912-932; Tm=55°C; amplification: 454bp). The PCR amplification of 559 and 454 nucleotides was checked on 1X agarose gel and amplicons were sequenced for further analyses (see below). For inconclusive samples, *i.e.* included in the uncertainly interval associated to the visualization method, a second PCR was performed using the same protocol as previously except the use of 2µl of RCA products. The samples still not conclusive after these two PCR reactions were then considered as negative for MSV. For all negative samples, PCR reactions amplifying the S1 locus area of chromosome 6 of rice were performed as internal control. Note that the amplicon size is used to discriminate the Asian rice *Oryza sativa* and the African rice *O. glaberrima* (935bp and 1384bp for *O. sativa* and *O. glaberrima*, respectively; Gnacadja et al. 2018).

Differences between percentage of MSV-positive fields or plants according to sites, rice production systems or years were assessed from contingency tables using Fisher’s exact test.

### Partial and complete genome sequencing

Amplicons obtained by detection PCR were sequenced by Sanger method (Genewiz-Azenta, South Plainfield, NJ, USA). We obtained usable sequences of varying sizes (from 491 to 550 nucleotides-long) subsequently used to identify which MSV strain is present at field or plant levels and to estimate the genetic diversity (see below) on the genetic fragment shared by all samples (*i.e.* 491nt, which correspond to 18.3% of the complete genome of MSV).

Two approaches were used to obtain complete genome sequences of MSV. First, the RCA products were digested with *BamH*I (New England Biolabs, Ipswich, MA, USA), inserted in pGEM-T Easy Vector (Promega, Madison, WI, USA) and cloned in JM109 competent cells (Promega) according to the manufacturer’s protocols. Alternatively, we performed two overlapping PCR on the RCA products with GoTaq Flexi (Promega) according to the manufacturer’s protocol to amplify the complete genome of MSV using primers targeting MSV-G (PCR#1: MSVg-2F and MSV-R559bp, Tm=57°C, amplification of 1956bp; PCR#2: MSV-F559bp and MSVg-2R: 5’-AGGCATGTCCGAACCGATGC-3’ at positions 980-999, Tm=57°C, amplification of 1811bp) and MSV-A (PCR#1: MSVg-2F and MSVa-3R: 5’-ATTGGCTCCAGCCTAACATCTTCC-3’ at positions 1898-1921, Tm=55°C, amplification of 1443bp; PCR#2: MSVa-1F: 5’-CGACGATGTAGAGGCTCTGCT-3’ at positions 1761-1781, and MSVa-1R, Tm=55°C, amplification of 1864bp). The obtained complete genomes sequences were deposited in GenBank (Accession Nos. from OR258386 to OR258402; Table Supp4). Two complete genome sequences of representative MSV isolates were used to obtain infectious clones (see below).

### Genetic diversity and phylogenetic analyses

Partial and complete genome sequences of MSV obtained during this study were compared to the 885 sequences described in (Kraberger et al. 2017). Multiple sequence alignments were performed using MUSCLE (Edgar 2004) implemented in SEAVIEW v4.7 (Gouy et al. 2010). Sequence pairwise identities of a selection of MSV sequences were calculated with SDT v1.2 (Muhire et al. 2014). Maximum likelihood phylogenetic trees were reconstructed with SEAVIEW using the best-fitted nucleotide substitution models (Tamura 3-parameter+G and GTR+G+I for the partial and the complete genome datasets, respectively) determined with MEGAX (Kumar et al. 2018) and 100 bootstrap replications. Phylogenetic trees were drawn using FigTree v1.3.1 (http://tree.bio.ed.ac.uk/software/figtree/). The genetic diversity of partial genome dataset was estimated using the best-fitted nucleotide substitution model, with standard errors of each measure based on 100 replicate bootstraps, as implemented in MEGAX.

Genetic differentiation of MSV populations according fields, sites, years and rice production systems were estimated by analysis of molecular variance (AMOVA) obtained by performing 1000 permutations as implemented in Arlequin v5.3.1.2 (Excoffier et al. 2005). Recombination signals in MSV from Burkina Faso were identified using the seven algorithms implemented in RDP v4.97 (Martin et al. 2015). Recombination events detected by at least 5 methods and with *P*-values below 10^-5^ were considered.

### Infectious clones and agroinoculations on rice

Three infectious clones of MSV-A and MSV-G were used during this study. First, the clone pBC-KS::MSV-A|R2| was obtained in previous studies and was identified to be highly pathogenic on maize (Isnard et al. 1998; Peterschmitt et al. 1996). Then, the clones pCAMBIA0380::MSV-A|53| and pCAMBIA0380::MSV-G|61| were built by gene synthesis (GENEWIZ-Azenta, South Plainfield, NJ, USA) of the genomic sequences obtained after PCR amplifications or cloning RCA products, respectively. More precisely, the MSV genomic sequences were synthetized based on the traditional technique of partial tandem repeats, *i.e.* with the highly conserved stem-loop region of ca. 50 nucleotides corresponding to the origin of replication repeated on both sides of the genomes (Urbino et al. 2008).

These infectious clones and the empty pCAMBIA0380 plasmid (as negative control, thereafter named pCAMBIA0380::Ø) were introduced into two strains of *Agrobacterium tumefaciens* (C58C1 and EHA105) by electroporation. Transformed *A. tumefaciens* colonies were plated and cultivated at 28°C on LB medium containing 50µg/ml of kanamycin and 25µg/ml of rifampicin (and 100µg/ml gentamycin for C58C1) and used for agroinoculation as described below after confirming the presence of inserts by PCR on colonies.

Agroinoculations were performed on two rice varieties representing the two cultivated rice species in Africa (*Oryza sativa indica* cv. IR64 and *O. glaberrima* cv. Tog5673) and one variety of maize (*Zea mays* cv. Golden Bantam). Plants were inoculated at the two-leaf stage (*i.e.* 7-10 days after seedlings) by pricking the apical meristem three times at different levels with the tip of 0.4mm needles previously dipped into agrobacterium colonies. Two independent assays have been performed. Both *A. tumefaciens* strains and pCAMBIA0380::Ø, pBC-KS::MSV-A|R2| and pCAMBIA0380::MSV-G|61| plasmids have been used for the first experiment. As no significant difference of infection was detected between the two *A. tumefaciens* strains (*X*²=1.101, *P*=0.294), we used only EHA105 for the second experiment. In addition, in this second experiment, the pathogenicity of the infectious clone pCAMBIA0380::MSV-A|53| was evaluated. For both experiments, symptom initiation and development have been monitored, and the number of leaves and the height of the inoculated plants were measured at 28 days post-inoculations (dpi). The fresh weight of inoculated plants was estimated only for the second experiment. Values were expressed for each treatment and cultivar in percentage according to negative controls.

Statistical analyses on these phenotypic measurements were performed using the software packages Statgraphics Centurion 15.1.02 (Stat Point technologies Inc., Warrenton, VA, USA). As the distribution of the plant size, number of leaves and fresh biomass was not normal according to Levene’s test of equality of error variances, we first analyzed our results by nonparametric (Kruskal-Wallis) test. However, as Kruskal-Wallis and ANOVA (parametric) tests gave similar results, and since ANOVA is robust to the partial violation of its assumptions and allows the analysis of factor interactions (post-hoc LSD, significance threshold at *P*<0.05) while Kruskal-Wallis does not, we also presented the results obtained with ANOVA analyses.

## RESULTS

### Identification of MSV in rice fields from Burkina Faso by viral metagenomics

Eighty-five rice samples were analyzed by a VANA-Illumina HiSeq approach, representing 43 and 42 rice fields surveyed in 2016 and 2017, respectively. The Illumina sequencing produced 288,971,924 raw sequences that were subsequently trimmed and corrected to extract a total of 1,459,838 contig sequences corresponding to 109.67 Mbases (Moubset et al., 2022). BlastN and BlastX analyses of these data showed that 1.9% of these contigs matched with viral sequences (project MP1 in Moubset et al., 2022) and notably those of maize streak virus detected in two fields surveyed in 2017. More precisely, 1490 MSV contigs were obtained from 17BM11 (98 contigs) and 17TG05 (1392 contigs) fields, including two long contigs from each field covering the complete genome of MSV. These long contigs respectively shared 99.01% and 98.92% homologies with the genomic sequence of a MSV isolate belonging to strain G collected in a wild *Poaceae* (*Urochloa lata*) from Nigeria (accession number EU628635.1). In addition to rice samples, six pools of wild plants, collected at the borders of one rice field per site, were analyzed by the same VANA-Illumina HiSeq approach (Table Supp1; https://doi.org/10.23708/1IPJAU). MSV was only identified in 17TG01w pool. The contig obtained from this sample covered the complete genome of this virus and Blast analyses revealed 99.37% homology with a MSV-G isolate from Nigeria collected in 2007 on the wild grass *Digitaria horizontalis* (accession number EU628634.1).

### Molecular detection of MSV in rice fields

As only two years of samplings were analyzed by VANA-Illumina and as the number of fields positive to MSV could be underestimated with this technique, we then opted for a targeted MSV detection strategy. For that, we used specific primers to detect by RCA-PCR the presence of this virus in rice fields: 43 fields in 2016, 40 in 2017, 49 in 2018 and 40 in 2019. Thus, we analyzed a total of 172 rice samples, each representing 16 plants from the same field collected the same year (Table Supp2).

A surprisingly high number (N=59) of rice samples were detected positive to MSV, suggesting that 34.3% of the surveyed fields were infected by this virus (Table 1). In addition, we noticed that MSV was detected in all localities and both production systems (*i.e.* irrigated and rainfed lowland fields), but with no significative difference for the 2016-2019 period (*X*²=6.424, *P*=0,267 and *X*²=0.170, *P*=0.680, respectively). However, the percentage of MSV-positive fields is significantly higher in 2018 and 2019 than in 2016 and 2017 (*X*²=44.269, *P*<0,001; Table 1).

**Table 1:**
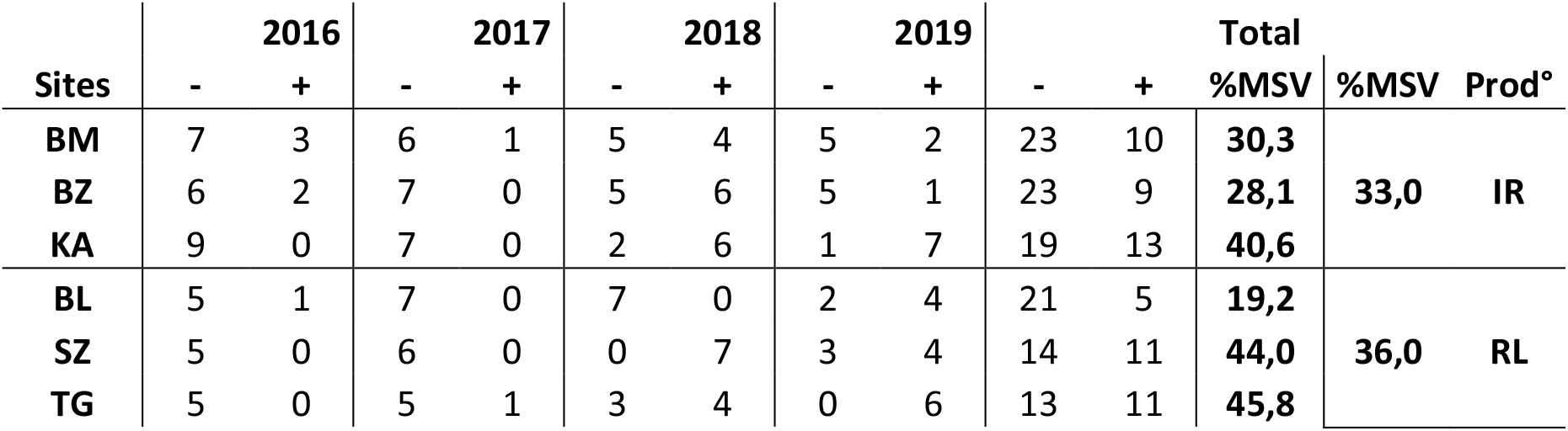
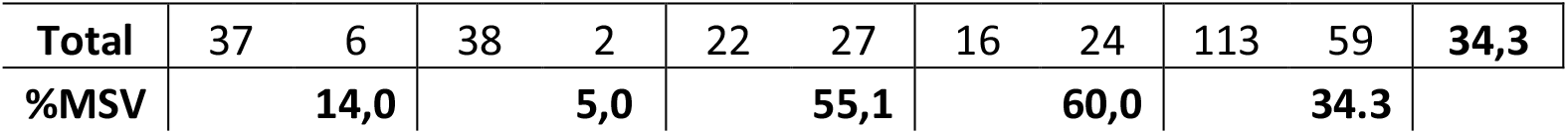
Detection of MSV in rice fields from 6 sites (BM: Bama; BZ: Banzon; KA: Karfiguela; BL: Badala; SZ: Senzon; TG: Tengrela) representing two rice production systems (IR: irrigated; RL: rainfed lowland) in 2016, 2017, 2018 and 2019. Percentage of MSV-positive fields are indicated (%MSV).

| Sites | 2016 |  | 2017 |  | 2018 |  | 2019 |  | Total |  | %MSV | %MSV | Prod° |
| --- | --- | --- | --- | --- | --- | --- | --- | --- | --- | --- | --- | --- | --- |
|  | - | + | - | + | - | + | - | + | - | + |  |  |  |
| **BM** | 7 | 3 | 6 | 1 | 5 | 4 | 5 | 2 | 23 | 10 | **30,3** | **33,0** | **IR** |
| **BZ** | 6 | 2 | 7 | 0 | 5 | 6 | 5 | 1 | 23 | 9 | **28,1** |  |  |
| **KA** | 9 | 0 | 7 | 0 | 2 | 6 | 1 | 7 | 19 | 13 | **40,6** |  |  |
| **BL** | 5 | 1 | 7 | 0 | 7 | 0 | 2 | 4 | 21 | 5 | **19,2** | **36,0** | **RL** |
| **SZ** | 5 | 0 | 6 | 0 | 0 | 7 | 3 | 4 | 14 | 11 | **44,0** |  |  |
| **TG** | 5 | 0 | 5 | 1 | 3 | 4 | 0 | 6 | 13 | 11 | **45,8** |  |  |

|  |  |  |  |  |  |  |  |  |  |  |  |
| --- | --- | --- | --- | --- | --- | --- | --- | --- | --- | --- | --- |
| **Total** | 37 | 6 | 38 | 2 | 22 | 27 | 16 | 24 | 113 | 59 | **34,3** |
| **%MSV** |  | **14,0** |  | **5,0** |  | **55,1** |  | **60,0** |  | **34.3** |  |

### Detection of MSV in wild and cultivated plants

In addition to rice samples, we first performed the MSV detection PCRs on the same pools of wild plants used with the VANA-Illumina approach (Table Supp1). As reported with the VANA-Illumina approach, MSV was detected in the 17TG01w. In addition, the detection PCR revealed that MSV was also present in 17TG02w. Both pool of samples combined several *Poaceae* plant species already described as host plants for MSV (as *Setaria palidefusca* Stapf et Hubb. and *Dactyloctenium aegyptium* Beauv.; Table Supp3; Kraberger et al., 2017).

Then, we analyzed individually symptomatic wild plants (including the wild rice species *Oryza longistaminata*) and cultivated *Poaceae* (maize and sugarcane) collected in 2016, 2017 and 2019 around and within the rice fields (Table Supp3). MSV was detected in a large proportion of samples: all maize (100%, N=8), most sugarcane (87.5%, N= 7) and a large fraction of wild plants (40% N=15), were identified MSV-positive (Table Supp3). More specifically, among the wild plant species identified, MSV was detected in *Digitaria horizontalis*, *Echinochloa colona*, *Eragrostis* sp. and *Oryza longistaminata*.

### Genetic diversity of MSV circulating in rice production areas

In order to inform/understand more/estimate about the MSV genetic diversity circulating in rice production areas from western Burkina Faso, we sequenced the amplicons obtained from molecular detection PCRs performed on rice, wild plants and cultivated *Poaceae* (maize and sugarcane). We obtained 29 sequences (Table Supp2) and phylogenetic reconstructions indicated that most of these sequences belonged to the strain G of MSV (27 out of 29, *i.e.* 93.1%), while only two corresponded to the strain A (16BZ09 and 18TG01, 2 out of 29, *i.e.* 6.9%; Table Supp2, Figure Supp2).

In addition to rice, the presence of MSV was also tested on wild grasses and cultivated *Poaceae* (maize and sugarcane) collected within or nearby the rice fields (Table Supp3). First, MSV-A was detected in 19 out of 30 (*i.e.* 63.3%) of plants showing symptoms (Table Supp3). Interestingly, MSV-G was only detected in one pool of symptomless wild grasses that were randomly collected from one site (17TG01w).

The genetic diversity estimated based on these fragments was low (0.0956 ± 0.0164 substitution/site in total, 0.0023 ± 0.0007 subst./site and 0.0028 ± 0.0017 subst./site for MSV-G and MSV-A, respectively). A genetic differentiation was observed between rice and *Poaceae* (*F_ST(Rice/Poaceae)_* = 0.760, *P*<0.001), which reflects the over-representation of MSV-G on rice and MSV-A on other *Poaceae*. However, taking each MSV strain separately, no genetic differentiation was revealed between the MSV-G or MSV-A isolates identified on rice or other *Poaceae* (*F_ST(Rice/Poaceae)_* < 0.001, *P*=1.000 and *F_ST(Rice/Poaceae)_* < 0.001, *P*=0.809 for MSV-G and MSV-A, respectively). In addition, genetic differentiation was never detected on MSV isolates according to rice production mode (All MSV: *F_ST(IR/RL)_* < 0.001, *P*=1.000; MSV-G: *F_ST(IR/RL)_* < 0.001, *P*=0.550; MSV-A: *F_ST(IR/RL)_* = 0.147, *P*=0.124). Similar results were obtained on other region of the genome (data not shown).

### Complete genomes of MSV

Based on the detection PCR and partial sequencing results, we selected several MSV-positive samples to obtain complete genomes sequences of MSV-A and MSV-G from different host species. Thus, we achieved 16 MSV complete genomes, 4 from rice fields, 6 from wild grasses, 2 from maize and 4 from sugarcane (Table Supp4). The sequence of 17BM11 MSV isolate (MSV-G|61|) was obtained after the cloning of RCA products. In all other cases, direct sequencing of PCR amplicons was performed. To these 16 sequences, we added the 3 complete genome sequences obtained by the VANA-Illumina approach (MSV-G on rice 17BM11|Contig1671| and 17TG05|Contig844|, MSV-G on wild grasses 17TG01|Contig711|) for further analyses (Table Supp4). Note that the MSV-G complete genome sequences obtained by Sanger and VANA-Illumina approaches on the same samples were highly similar (11 and 1 variable sites for 17BM11 and 17TG01w, *i.e.* 99.6% and 100.0% genetic identity, respectively).

The 19 sequences obtained during this study were compared to MSV complete genome dataset gathering 885 sequences from all the MSV strains, including 8 MSV-G sequences from West Africa (Nigeria and Mali) and Gran Canaria on wild grasses and 695 MSV-A sequences from all over Africa with 5 MSV-A collected in 2008 in Burkina Faso on maize (Kraberger et al. 2017). Analyses to detect recombination events on this sequence dataset demonstrated that none of the 19 complete genome sequences obtained during this study were recombinant. Phylogenetic analyses showed that the MSV-G isolates identified during this study were closely genetically related (more than 98.9% of genetic identity; Figure 2). As previously demonstrated with partial genome analyses, no genetic differentiation was observed between MSV collected in rice or other *Poaceae* (*F_ST(Rice/Poaceae)_* = 0.362, *P* < 0.001) or between rice production mode (*F_ST(IR/RL)_* < 0.001, *P* = 1.000). MSV-G isolates from western Burkina Faso were genetically related to those identified in Nigeria in 2007 and Mali in 1987 (more than 99% of identity with EU628632.1 and EU628634.1). Similarly, we noted that the MSV-A isolates identified in this study belong to a clade of isolates exclusively identified in West Africa and were closely genetically related (more than 99.4% of identity), with no genetic differentiation with those collected in rice or other *Poaceae* (*F_ST(Rice/Poaceae)_* < 0.001, *P* = 0.985). Surprisingly, these isolates were more genetically related to MSV-A isolates collected in Nigeria in 2015 (between 99.5% and 99.9% of genetic identity with KX787926.1 and KX787927.1 for instance, ca. 1250km distance between these isolates and those from our study area) than with MSV-A identified in Burkina Faso in 2008 on maize (between 96.9% and 96.6% of genetic identity, ca. 300km distance between isolates; Figure 2).

**Figure 2:**
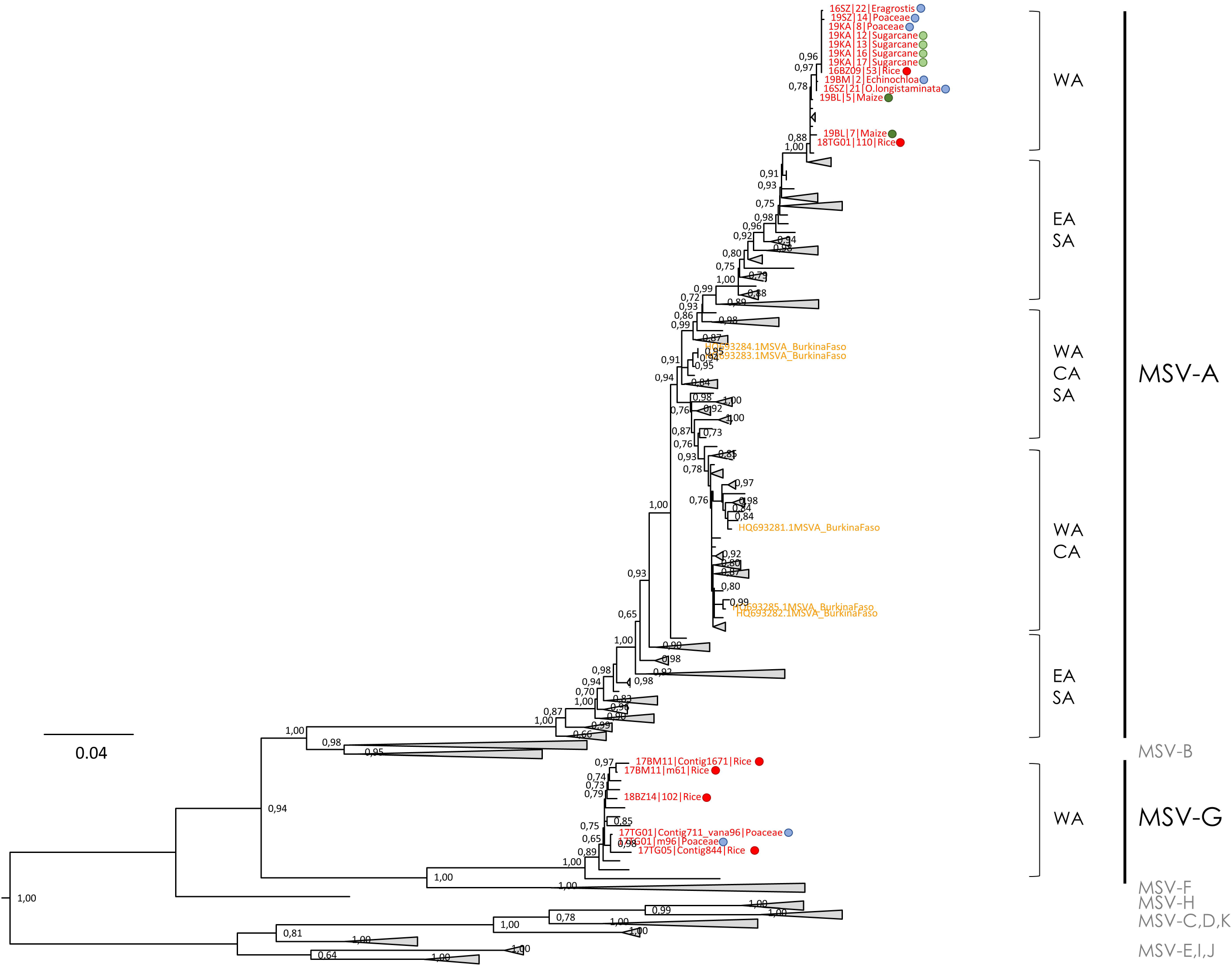
Condensed phylogenetic tree reconstructed by maximum-likelihood based on complete MSV genome sequences from western Burkina Faso obtained during this study (names in red) and available from public database (in orange for those from Burkina Faso). Numbers at each node correspond to bootstrap values based on 100 replicates (only values above 0.70 are reported). The host plants from which these sequences were identified are indicated by the colored circles (red: rice; dark green: maize; light green: sugarcane; blue: wild grasses). The clades corresponding to MSV strains (from A to K) and the geographical origin (CA: Central Africa; EA: East Africa; SA: Southern Africa; WA: West Africa) of the sequences are mentioned.

### MSV prevalence in rice fields

To assess MSV prevalence in rice fields, we analyzed individually the 16 plants of 12 rice fields previously identified as MSV-positive in 2018 by detection PCR (Table Supp2). These fields have been selected to represent different sites (BM, BZ, SZ, KA and TG), genetic diversity of MSV in rice fields (MSV-G *vs*. MSV-A) and rice production mode (irrigated *vs*. rainfed lowland). Badala (BL) site was not included in this analysis as no MSV-positive field has been identified (Table Supp2).

We noticed that MSV-G was frequently detected (32.1% of the plants), with prevalence varying between 25.0% and 62.5% according to the field (Figure 3, Figure Supp3, Table Supp5) but with no significant difference between fields (X² = 16.167, *P* = 0.135). Interestingly, no significant difference was also revealed between production system (X² = 0.185, *P* = 0.667).

**Figure 3:**
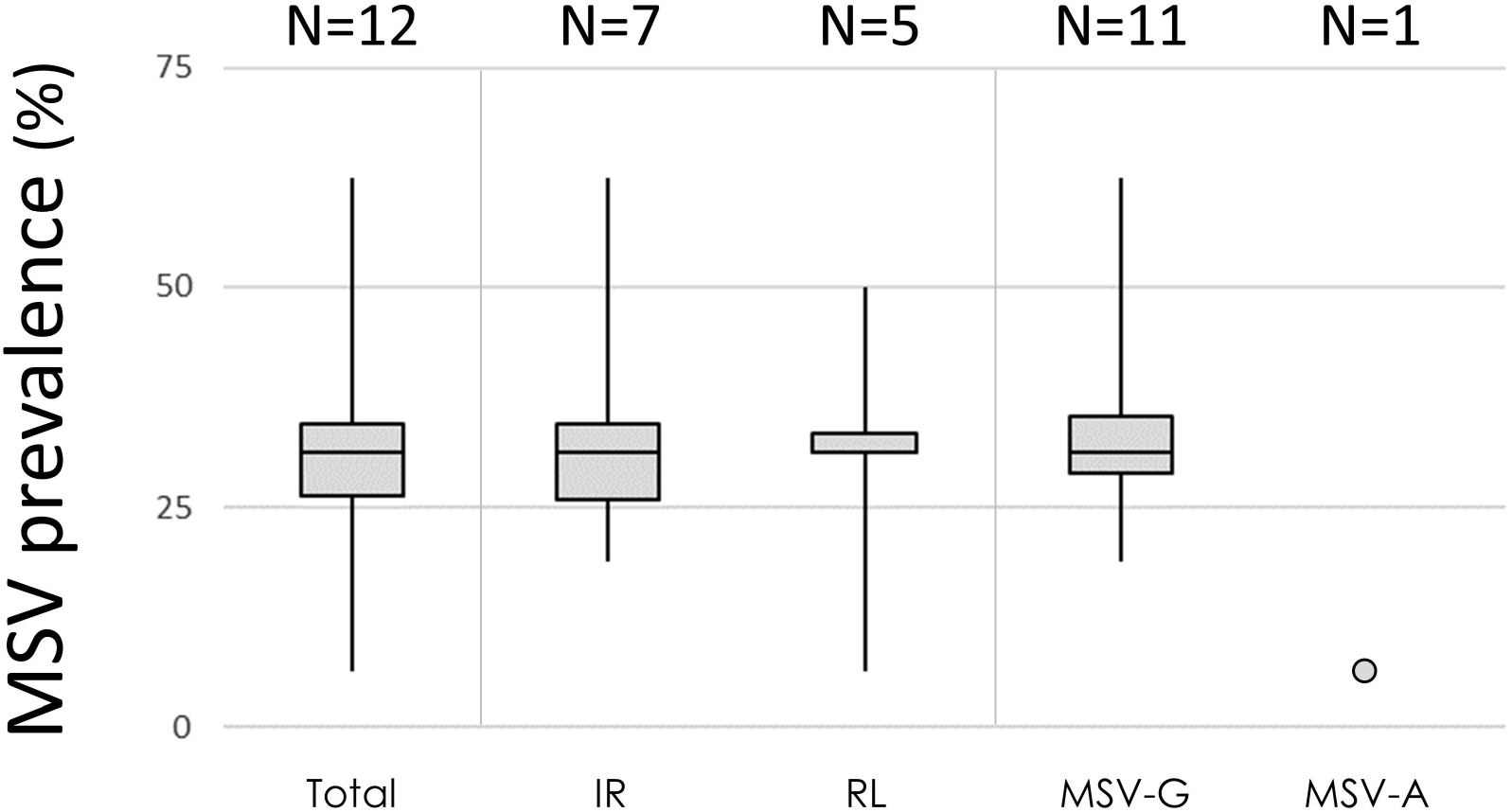
Prevalence estimation of MSV-G and MSV-A in 12 rice fields representing two rice production systems (IR: irrigated areas; RL: rainfed lowlands) based on the 15-16 individual plants in each field collected without *a priori* over a grid in 2018. The numbers of rice fields are indicated.

The genetic diversity estimated for MSV-G circulating within 18BM13, 18BZ14, 18SZ06 and 18KA08 with the detection PCR fragments was low (0.0058±0.0048 subst./site, 0.0066±0.0035 subst./site, 0.0060±0.0059 subst./site and 0.0035±0.0025 subst./site, respectively), and no genetic differentiation between fields have been observed (*F_ST_*_fields_<0.001, *P*=1.000).

For MSV-A, in addition to not being frequently detected in rice fields, the prevalence of this strain estimated in 18TG01 field was drastically lower than those of MSV-G (6.3%; Figure 3, Figure Supp3). No coinfection between MSV-G and MSV-A was observed during this study.

### Experimental validation of MSV pathogenicity on rice

We performed two independent experiments to test the pathogenicity of three MSV infectious clones: two obtained from the isolates identified during this study in rice plants from Burkina Faso (pCAMBIA::MSV-G|61| and pCAMBIA::MSV-A|53|) and one from an isolate identified in maize plant from Reunion Island (pBC-KS::MSV-A|R2|; Figure Supp4). For both experiments, we observed the emergence of light streaks on leaves for some rice plants after 14 days post-agroinoculation (dpi) of the infectious clones, in *Oryza sativa indica* cv. IR64 and *O. glaberrima* cv. Tog5673. Then, these symptoms drastically evolved into clear and marked streaks, both on the leaves where the streaks were first observed and on the systemic and emergent leaves until 28 dpi, followed with drastic reduction of plant growth (Figure 4). Correlation between symptoms and MSV accumulation in systemic leaves was validated by PCR and Sanger sequencing (data not shown).

**Figure 4:**
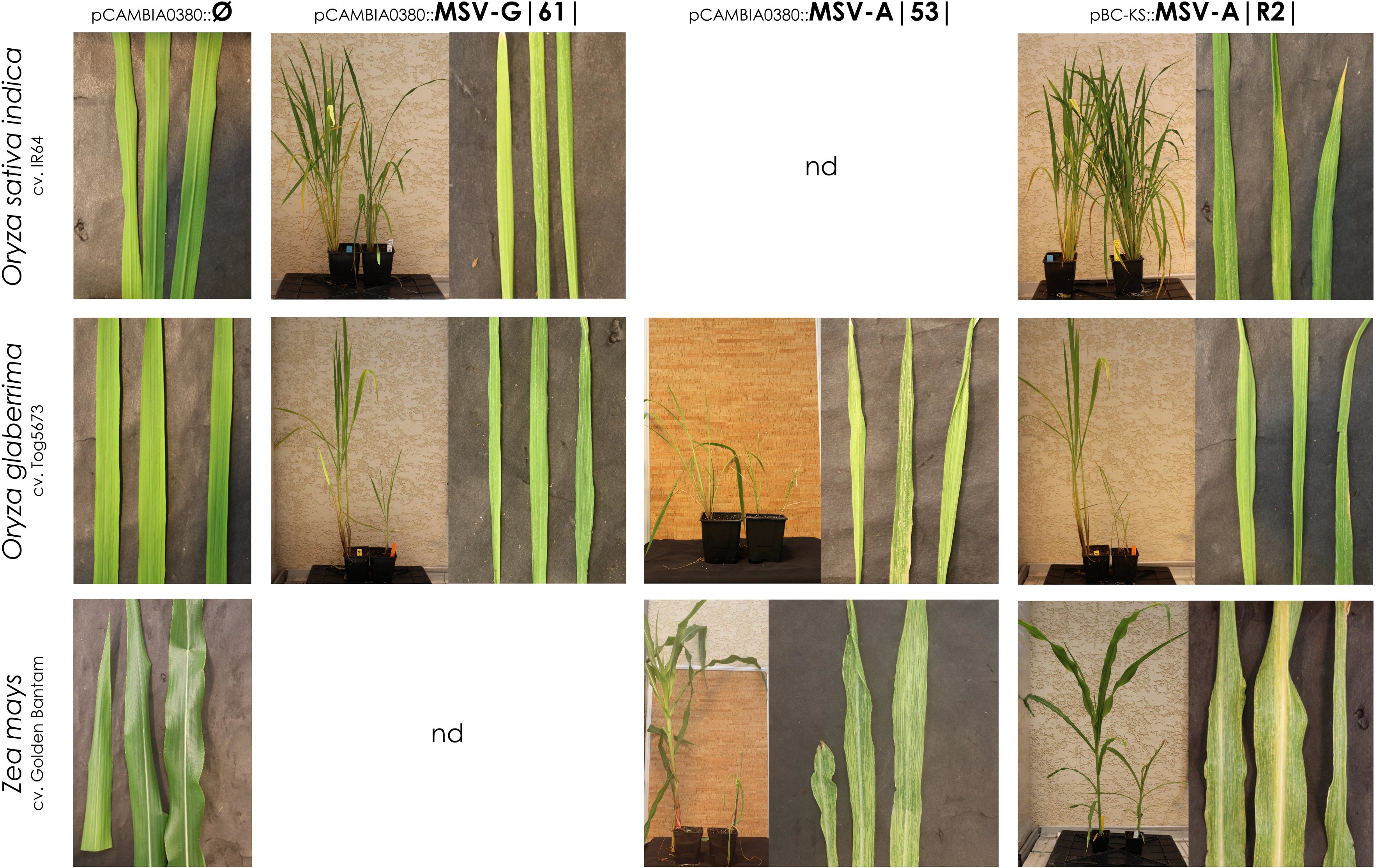
Symptom observation in rice (*Oryza sativa indica* cv. IR64 and *O. glaberrima* cv. Tog5673) and maize (*Zea mays* cv. Goldem Bantam) associated to MSV infection after agroinoculations of pCAMBIA::MSV-G|61|, pCAMBIA::MSV-A|53| and pBC-KS::MSV-A|R2| infectious clones. Photos of the whole plants showing a negative control plant (left) and infected plants (right) were taken 60 days post-inoculation (at 28dpi only for pCAMBIA::MSV-A|53| on *O. glaberrima* cv. Tog5673), those of systemic and emergent leaves at 28dpi. nd: not determined because no MSV infected plant was identified.

Unfortunately, most likely due to the versatility of the agroinoculation process on rice, only few cases of successful infections have been observed, and not all agroinoculation modalities (*i.e.* infectious clone *vs*. plant species) succeed during the same experiment. Nevertheless, by clumping the results of the two experiments, the MSV agroinoculation led to successful infection in 46 out of 770 plants (*i.e.* 6.0%; Table 2, Table Supp6). No significant difference of MSV transmission efficiency was observed between the three infectious clones on *O. sativa indica* cv. IR64 (X²=2.144, *P*=0.342) and *O. glaberrima* cv. Tog5673 (X²=0.764, *P*=0.682). However, MSV was more efficiently transmitted to Tog5673 than to IR64 (42 out of 392 plants, *i.e.* 10.7% and 4 out of 378 plants, *i.e.* 1.1% for Tog5673 and IR64, respectively; X²=31.943; *P*<0.001), and so whatever the infectious clone (X²=12.968, *P*<0.001; X²=17.216, *P*<0.001 and X²=2.856, *P*=0.091 for pCAMBIA::MSV-G|61|, pCAMBIA::MSV-A|53| and pBC-KS::MSV-A|R2|, respectively; Table 2, Table Supp6).

**Table 2:** Total number of non-symptomatic (-) and symptomatic (+) plants observed 28 days post-inoculation (dpi) of pCAMBIA0380::Ø (negative control), pCAMBIA0380::MSV-G|61|, pCAMBIA0380::MSV-A|53| and pBC-KS::MSV-A|R2| in *Oryza sativa indica* cv. IR64, *O. glaberrima* cv. Tog5673 and *Zea mays* cv. Golden Bantam. Percentages of symptomatic plants are indicated with %(+).

| Infectious clone | *O. sativa indica* cv. IR64 |  |  | *O. glaberrima* cv. Tog5673 |  |  | *Zea mays* cv. Golden Bantam |  |  |
| --- | --- | --- | --- | --- | --- | --- | --- | --- | --- |
|  | - | + | %(+) | - | + | %(+) | - | + | %(+) |
| ∅ | 16 | 0 | 0.0 | 20 | 0 | 0.0 | 3 | 0 | 0.0 |
| pCAMBIA0380::MSV-G\|61\| | 198 | 3 | 1.5 | 185 | 20 | 9.8 | 84 | 0 | 0.0 |
| pCAMBIA0380::MSV-A\|53\| | 127 | 0 | 0.0 | 110 | 16 | 12.7 | 67 | 1 | 1.5 |
| pBC-KS::MSV-A\|R2\| | 49 | 1 | 2.0 | 55 | 6 | 9.8 | 4 | 1 | 20.0 |
| **Total MSV-G/A** | **374** | **4** | **1.1** | **350** | **42** | **10.7** | **155** | **2** | **1.3** |

Twenty-eight days after agroinoculation, we measured the height and the number of leaves of MSV-infected and non-infected plants. We noticed that the earlier the emergence of symptoms, the greater was the impact on plant growth, leading to high between-plant heterogeneity in the dataset. While MSV infection did not systematically affect the number of leaves per plant (Kruskal-Wallis: *F*=32.706, *P*<0.001; ANOVA: *F*_6,150_=7.750, *P*<0.001; Figure Supp5A), we showed that the height of plants was significantly reduced by MSV infection (Kruskal-Wallis: *F*=68.751, *P*<0.001; ANOVA: *F*_6,150_=22.240, *P*<0.001; Figure 5). In addition, the evaluation of the plant fresh weight during the second experiment suggests that MSV infection has an impact on the biomass production in rice (Kruskal-Wallis: *F*=30.877, *P*<0.001; ANOVA: *F*_4,122_=4.850, *P*=0.001; Figure Supp5B).

**Figure 5:**
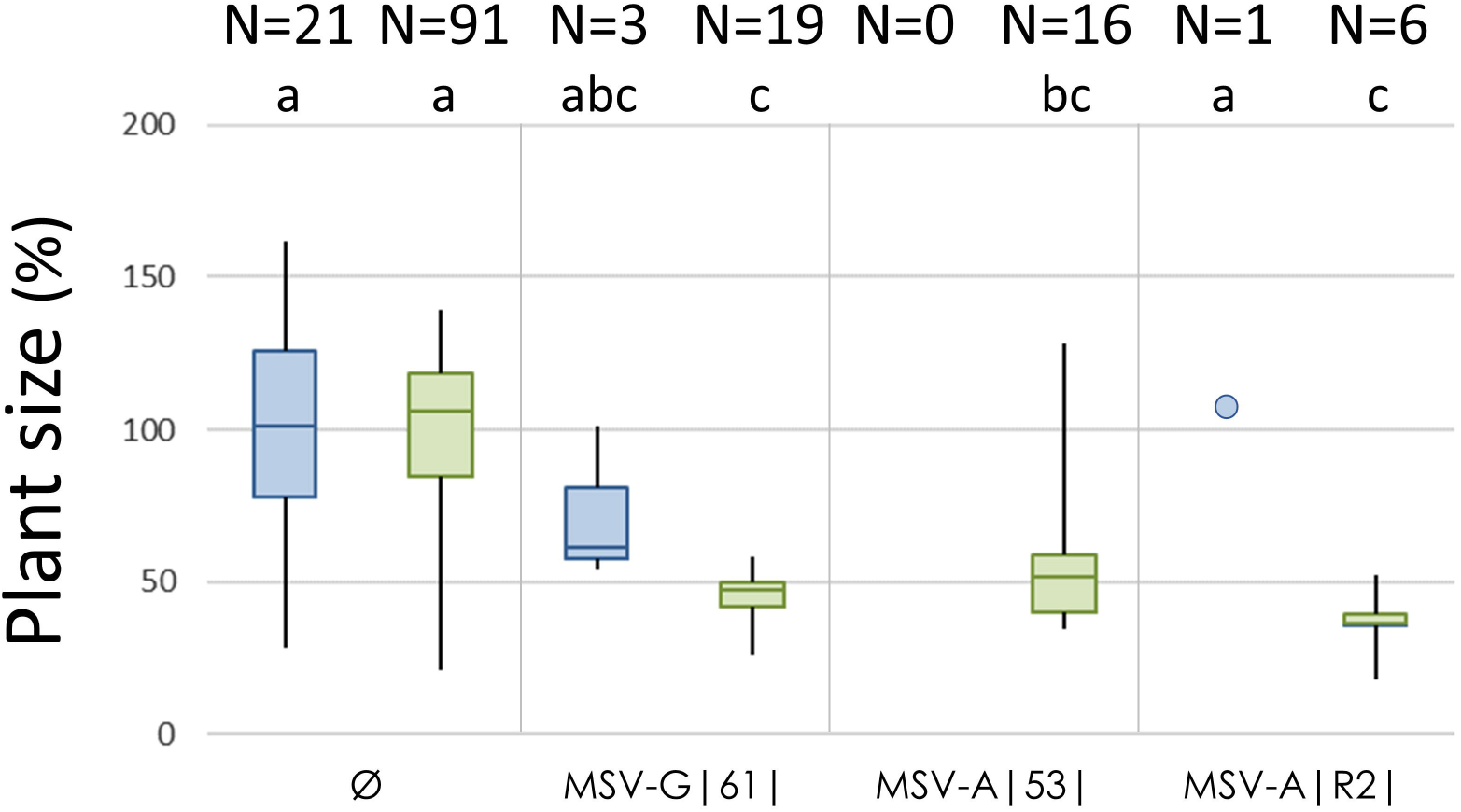
Maximal height of MSV infected plants (inoculated with pCAMBIA0380::MSV-G|61|, pCAMBIA0380::MSV-A|53| and pBC-KS::MSV-A|R2|) normalized according to negative control plants (inoculated with pCAMBIA0380::Ø) for *O. sativa indica* cv. IR64 (blue) and *O. glaberrima* cv. Tog5673 (green). The numbers of plants used to obtain these data are indicated. The statistically identical groups are mentioned (a,b,c). The value obtained with the unique IR64 infected after pCAMBIA::MSV-A|R2| agroinoculations are reported with a blue point.

In parallel, the infectious clone pBC-KS::MSV-A|R2| was agroinoculated to maize as positive control, and one symptomatic and MSV infected plant was obtained (1 out of 5 plants; Table 2, Figure 4). The emergence of typical symptom of MSV on maize was observed after 9 dpi and MSV-A|R2| accumulation was validated by PCR and Sanger sequencing (data not shown). By contrast, only one maize plant out of 68 (*i.e.* 1.5%) agroinoculated with pCAMBIA0380::MSV-A|53| started to develop typical symptoms of MSV after 21 dpi (Table 2; Table Supp6). Successful infection was never obtained on maize with pCAMBIA0380::MSV-G|61| agroinoculation (Table 2; Table Supp6).

### Observation in fields of symptoms putatively associated to MSV-infection

As phenotypes and symptoms of the collected plants in rice fields and pictures of plants have been recorded during the sampling process, we tried *a posteriori* to associate the results of MSV detection with the presence of the symptoms that could be associated to MSV infection (based on the results of the experimental MSV agroinoculations). We first noticed that “white stripes” were frequently observed in rice plants in 2018 and 2019, whereas it was never reported in 2016 and 2017 (Table Supp2), which could parallel with the higher frequency of MSV-positive fields on the last two years of the survey.

However, no significant association between “white stripes” phenotypes and MSV-positive samples have been observed at the field level (X²=1.442, *P*=0.230 for the 2016-2019 period; X²=1.199, *P*=0.274 and X²=2.063, *P*=0.151 for 2018 and 2019, respectively), or at the individual plant level (based on plant samples from 12 fields in 2018 used to estimate MSV prevalence: X²=0.919, *P*=0.340). Only a significant association between symptoms and MSV-positive samples was noticed in the site Badala at the field level (BL: *X*²=8.864, *P*=0.003). Nevertheless, we identified 4 symptomatic rice plants in fields that we confirmed *a posteriori* to be infected by MSV-G (detection PCR and Sanger sequencing; cf Figure Supp2).

## DISCUSSION

By combining epidemiological surveys in fields (Barro et al. 2021a), viral metagenomics approach (VANA-Illumina; Moubset et al. 2022), molecular epidemiology and experimental infections, we described for the first time the epidemiology, the genetic diversity and the pathogenicity of MSV that had never been reported before on rice (Table 1). Indeed, although MSV has a very large host range infecting dozens of plant species of the *Poaceae* family (Kraberger et al. 2017) and demonstrated to be transmitted experimentally on rice by leafhoppers (Damsteegt 1983; Konaté and Traoré 1992), no study had shown until now the extent of MSV epidemics in rice fields from Africa.

Based on rice and wild grasses samples collected in 2016 and 2017, sequences sharing identity with MSV strain G were identified from two pools of rice plants and one pool of wild plants using the VANA-Illumina approach. Although the VANA-Illumina approach is extremely valuable for estimating the diversity of virus populations and for expanding the knowledge about virus species circulating within the environment (Moubset et al. 2022), the comparison of MSV detection by VANA-Illumina and RCA-PCR suggested that the detection threshold was lower with RCA-PCR than VANA-Illumina. Indeed, while MSV was detected by both methods for one rice field; 7 additional rice fields were identified positive to MSV in 2016 and 2017 by RCA-PCR (Table Supp2). Conversely, we noticed that one field was detected positive to MSV by VANA-Illumina approach but not by RCA-PCR. As the plant material used for these two approaches were not strictly identical (independent virion or nucleic acid extractions), we can assume that we could generate some discrepancy between extractions and so MSV detection if the virus is not uniformly distributed within the infected plants. In addition, analysis by pool of several plants together could also contribute to weaken our ability to consistently detect MSV in rice or wild grass samples. Altogether, we can consider that the percentage of rice fields where MSV was circulating was potentially underestimated in this study.

Despite all these putative biases of detection, MSV was surprisingly frequent in rice landscape of western Burkina Faso, especially in 2018 and 2019 (Table 1, Table Supp2). Indeed, the frequency of MSV-positive fields was similar to the frequency of symptoms caused by another well-known virus, the rice yellow mottle virus (RYMV, *Solemoviridae*), in these same fields and during the same time period (34.3% and 30.2% for MSV and RYMV, respectively Barro et al. 2021a). In addition, similarly to RYMV, the production system (irrigated *vs*. rainfed lowland) does not have a significant effect on the percentage of MSV-positive fields (Table 1). However, we noticed a significant variation of the frequency of MSV-positive fields between years (Table 1), which could be due to variations in climatic conditions and insect vector populations (as suggested for MSV epidemiology in maize cropping areas from Reunion Island; Reynaud et al. 2009).

Two MSV strains were identified in rice fields (MSV-G and MSV-A) and MSV-G was drastically more frequent than MSV-A (Figure Supp2, Table Supp2). As far as we know, MSV-G has only been identified on wild grasses, whereas MSV-A has been reported both on wild grasses and cultivated *Poaceae*, such as maize and sugarcane (Kraberger et al. 2017). However, while specific primers were used in this study to detect MSV-G and MSV-A by (RCA-)PCR, the genuine diversity of MSV strains in rice agroecosystems could have been underestimated even if no other MSV strain has been identified by metagenomic VANA-Illumina approach.

During this study, MSV-A was only identified in 2 fields (Table Supp2) for which rice and maize were cultivated alongside or in rotation (data not shown). Interestingly, no genetic differentiation between the MSV isolates collected in rice and other wild or cultivated *Poaceae* growing around the rice fields was detected, suggesting that MSV-G and MSV-A circulate indifferently between these host plant species. Nevertheless, we noticed in our study that MSV-G was only detected in symptomless wild *Poaceae* that were randomly sampled while MSV-A was only identified on plants specifically collected because presenting symptoms of viral infection (Table Supp3). These results suggest that MSV-A could be more aggressive, *i.e.* inducing more symptoms, than MSV-G. In parallel, these results imply that the prevalence and the spatial distribution of MSV-G could be underestimated because of probably more limited symptom induction compared to MSV-A and so a lower propensity to be collected during plant pathology studies, especially within the wild compartment (Lefeuvre et al. 2019).

Based on 12 MSV-positive rice fields surveyed in 2018, we estimated that MSV prevalence within these fields was 32.1% on average and that MSV-A was overall less prevalent than MSV-G (6.3% *vs*. 18.8-62.5%; Figure 3, Table Supp5). Note that, as previously discussed, the high threshold of MSV detection at the field level (*i.e.* pool of 16 plant leaves from each field) could imply that only fields with high proportion of MSV infected plants were detected positive by RCA-PCR, and so that the average prevalence could be overestimated. Nevertheless, the analysis of these 12 fields revealed the impressive MSV prevalence in rice fields from Burkina Faso in 2018, with no significant difference between sites or rice production system. These results suggested that MSV can circulate efficiently between these agri-environment, probably because of the flight performance of the insect vector (Asanzi et al. 1995) and the absence of varietal differentiation between irrigated and rainfed lowland rice fields in this area (Barro et al. 2021b).

We validated in controlled conditions by agroinoculation the pathogenicity of MSV-G and MSV-A isolates. Both MSV strains are able to infect both rice species cultivated in Africa (*Oryza sativa* and *O. glaberrima*). Although the percentage of successful infection was limited (1.1% and 10.7% on *O. sativa* cv. IR64 and *O. glaberrima* cv. Tog5673; Table 2, Table Supp6), MSV infection induced severe symptoms (Figure 4) generally associated with a significant reduction of the size (Figure 5) and biomass production (Figure Supp5B) of infected rice plants. The efficiency of agroinoculation leading to infection was similar between MSV-G and MSV-A on each rice variety/species (Table 2). However, the percentage of successful infection of MSV-G, MSV-A or both strains combined was significantly higher in *O. glaberrima* cv. Tog5673 than *O. sativa indica* cv. IR64 (Table 2). Thus, this result suggests that, despite the restricted efficiency of agroinoculation method to transmit the virus in rice or the restricted number of rice varieties used in this study, the fitness of MSV could be higher on *O. glaberrima* than *O. sativa*. Further studies including more cultivars of each rice species and insect vectors will be required to confirm this hypothesis.

Interestingly, similar symptoms to those induced by experimental MSV agroinoculation have been observed in rice fields (Table Supp2, Table Supp5). However, no *a posteriori* significant association has been identified between these symptoms and MSV detection by PCR, at field or plant levels. This lack of association could be related to 1) the low detection threshold of MSV by PCR at the field level (cf. above), and/or 2) the rice varieties used in Burkina Faso as well as the environmental conditions that could reduce the intensity of symptoms in fields, and/or 3) to the misinterpretation of specific symptoms of MSV infection and the confusion with symptoms induced by other viral infections circulating in these area (Barro et al. 2021a; Sereme et al. 2014). Nevertheless, association between symptoms, MSV detection and MSV sequencing have been validated with 4 plants that have been specifically collected because showing symptoms that could be attributed to viral infection (cf. Figure Supp2). Note that the BD10 PCR amplifications, used as an internal control of the DNA extraction quality (cf. Material & Methods), showed that all the rice samples collected in Burkina Faso for this study exclusively correspond to *O. sativa* varieties, which is concordant with previous results from the same area (Barro et al. 2021b).

In future, the use of quick and sensitive diagnostic tool for detection of MSV directly in rice field, like loop-mediated isothermal amplification (LAMP) assays (Tembo et al. 2020), will help to unraveled the association between MSV infection and symptoms in rice, and will contribute to better know the MSV epidemiology in rice fields from Africa.

The MSV agroinoculation on maize led to successful infection for pBC-KS::MSV-A|R2| and pCAMBIA0380::MSV-A|53| infectious clones. The isolate MSV-A|R2|, collected in the Reunion Island and cloned after serial passages on almost resistant inbred maize lines, was defined to be one of the most pathogenic isolates analyzed in Isnard et al. 1998). In comparison, the isolate MSV-A|53|, collected in rice from Burkina Faso (16BZ09; Table Supp2), was also able to infect maize. These results demonstrate the ability of these isolates to infect both maize and rice whatever their host plant of origin, their geographical areas of origin and their genetic divergence (Figure Supp4).

Interestingly, we never detected MSV-G infection on maize during our agroinoculation assays (84 plants in total; Table 2). If this trend is confirmed (especially by assays involving insect vectors in order to increase MSV transmission efficiency), the asymmetric pattern of infection between MSV-G and MSV-A on rice and maize could shed light on the evolutionary history and host adaptation of MSV. Indeed, MSV-G has so far only been identified in wild grasses from West Africa while intensive analyses of MSV diversity in cultivated *Poaceae* as maize, sorghum and sugarcane have been performed (Kraberger et al. 2017). The results obtained in our study suggest the role of rice fields as boosters of MSV-G epidemics in the environment because of the homogeneity, density and spatial distribution of this crop (Anderson et al. 2004). Thus, as rice and maize are frequently grown within the same fields or areas in Africa, rice could have played an indirect role on the MSV adaptation to maize by increasing the probability of co-infections between MSV-G and MSV-B which subsequently led to the emergence of MSV-A by recombination (Harkins et al. 2009).

Altogether, in addition to fulfill the Koch’s postulates by experimental MSV inoculations on two rice species cultivated in Africa, this study suggests that MSV could be a significant pathogen for rice cultivation in Africa as it was found highly prevalent in rice fields from Burkina Faso and induced severe symptoms on rice plants in controlled conditions. Actually, MSV-G was, up to now, detected in wild grasses from West Africa including Nigeria and Mali. As these two countries correspond to the most important rice producer countries in West Africa (FAOSTAT 2022, https://data.un.org/Data.aspx?d=FAO&f=itemCode%3A27), we can assume that MSV-G could be epidemic in these countries. In addition, we experimentally confirmed that two MSV-A isolates genetically distinct and collected on rice in West Africa or maize in East Africa are able to infect efficiently both rice species (Figure 4, Figure Supp4). Thus, as MSV-A was reported almost everywhere in Africa (Kraberger et al. 2017) where rice is produced at large scale (for instance, West Africa: Benin, Burkina Faso, Ghana, Nigeria; Central Africa: Cameroon, Central African Republic, Chad; East Africa: Ethiopia, Kenya, Madagascar, Tanzania, Uganda), our results suggest lurking MSV epidemics on rice across the continent and promote constant and updated epidemiological surveillance of rice production in Africa.

## ACKNOWLEDGEMENT

This work was performed thanks to the facilities of the “International joint Laboratory LMI PathoBios: Observatory of plant pathogens in West Africa: biodiversity and biosafety” (www.pathobios.com; twitter.com/PathoBios). We are very grateful to Sylvain Zougrana, Yacouba Kone, Edouard Kabore, Moustapha Koala, Manaka Douanio, Amadou Diallo, Fabrice Nikiema, Bouda Zakaria, Ouedraogo Alassane, Fatoumata Gnacko, Daouda Hema, Drissa Tou, Traoré Momouni, Dabire Roméo, Martial Kabore and Noël Ouattara for their contributions to the fieldwork in Burkina Faso. We thank the rice farmers from Badala, Bama, Senzon, Banzon, Tengrela and Karfiguela for their kind collaboration. We also thank Michel Peterschmitt, Cica Urbino and Martine Granier for the valuable discussions and for kindly providing us the infectious clone pBC-KS::MSV-A(R2), and Jamel Aribi for plant care in IRD greenhouses.

## CAPTIONS

**Figure Supp.1:**
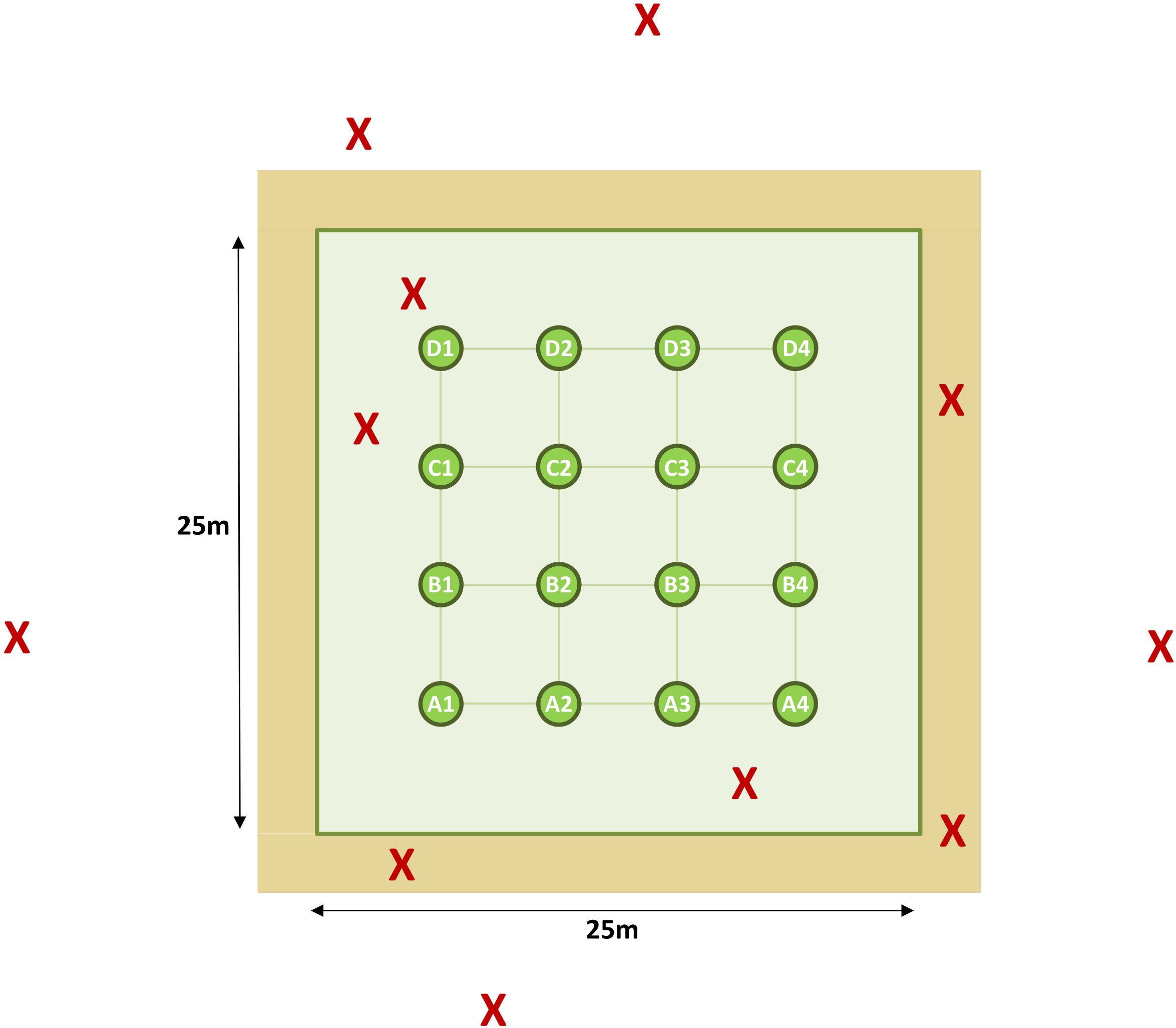
Schematic representation of the sampling protocol. The green circles labelled from A1 to D4 represent the 16 rice plants regularly collected over a grid without *a priori*, *i.e.* not based on their symptomatic status, in a rice field (green square). Wild grasses were collected without *a priori* on the borders of the rice field (light brown area) and additional plants (rice, maize, sugarcane or wild grasses; red crosses) presenting symptoms that could be related to virus infection collected within or nearby the rice field.

**Figure Supp.2:**
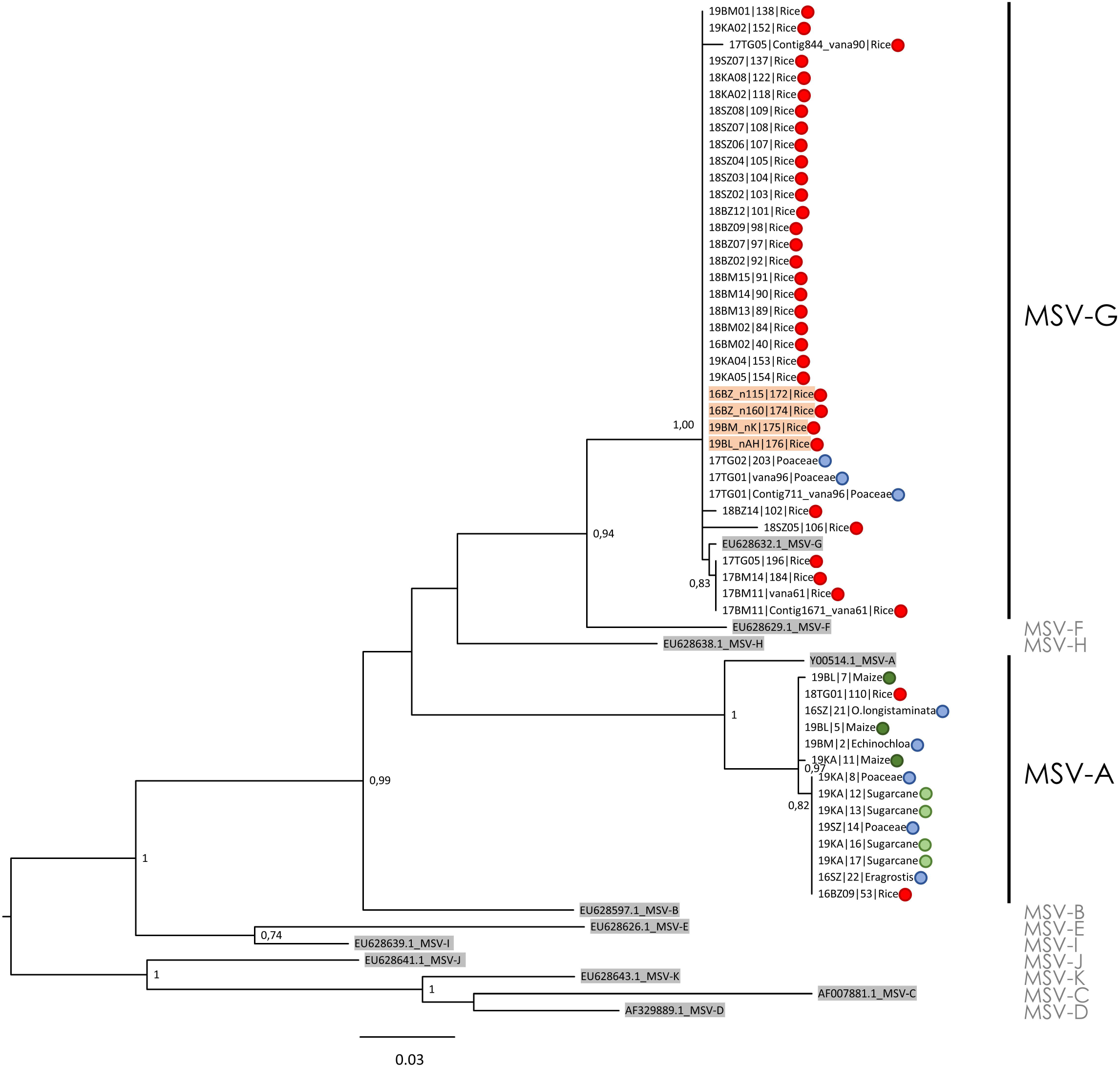
Phylogenetic tree reconstructed by maximum-likelihood based on partial MSV genome sequences (491 nucleotides long) of reference (highlighted in grey) and obtained with RCA-PCR (primers MSV-F559pb and MSV-R559pb) for MSV detection in rice (red points) or other *Poaceae* (dark green: maize; light green: sugarcane; blue: wild grasses). Sequences obtained from symptomatic rice plants are highlighted in red. Numbers at each node correspond to bootstrap values based on 100 replicates (only values above 0.70 are reported). The clades corresponding to MSV strains (from A to K) are indicated.

**Figure Supp.3:**
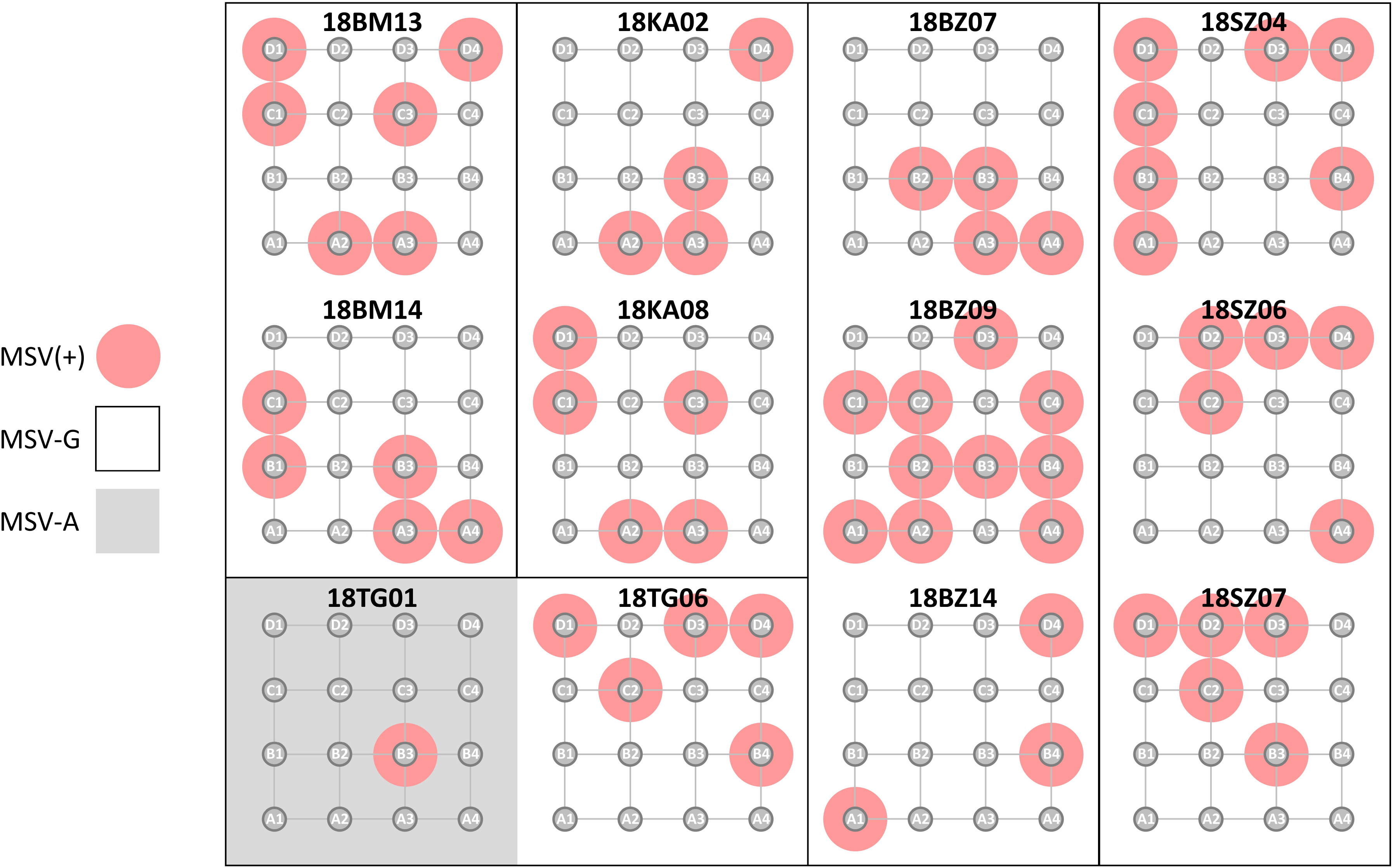
Schematic representation of the spatial distribution of the MSV-positive rice plants determined in 12 fields surveyed in 2018. The 16 rice plants regularly collected without *a priori* over a grid correspond to the grey circles labelled from A1 to D4, the red highlights show the MSV-positive ones. The rice field where MSV-A was detected is highlighted in grey.

**Figure Supp.4:**
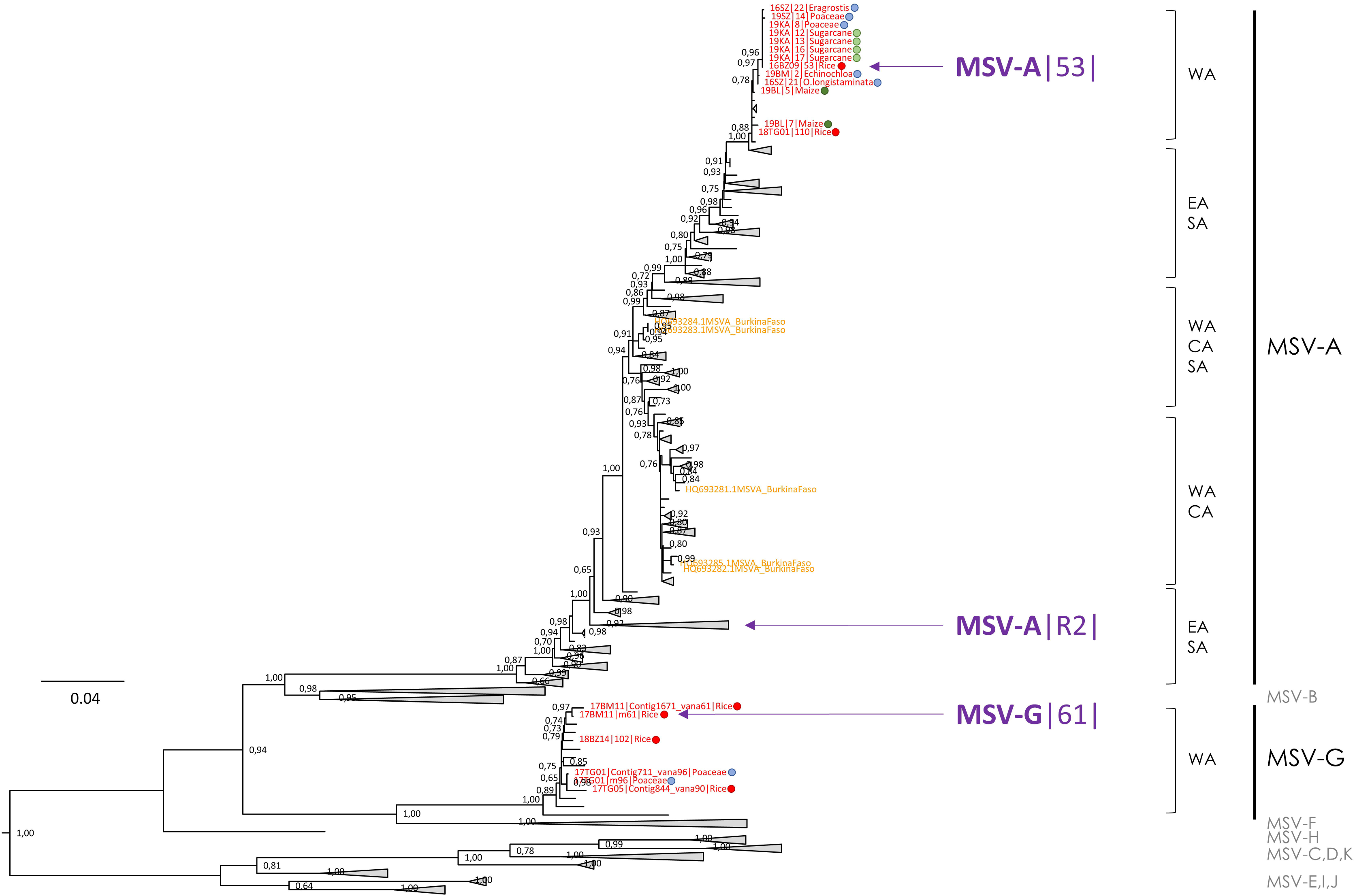
MSV sequences used to build infectious clones (MSV-A|53|, MSV-A|R2|, MSV-G|61|) located in the condensed phylogenetic tree reconstructed by maximum-likelihood based on complete MSV genome sequences from western Burkina Faso obtained during this study (names in red) and available from public database (in orange for those from Burkina Faso). Numbers at each node correspond to bootstrap values based on 100 replicates (only values above 0.70 are reported). The host plants from which these sequences were identified are indicated by the colored circles (red: rice; dark green: maize; light green: sugarcane; blue: wild grasses). The clades corresponding to MSV strains (from A to K) and the geographical origin (CA: Central Africa; EA: East Africa; SA: Southern Africa; WA: West Africa) of the sequences are mentioned.

**Figure Supp.5:**
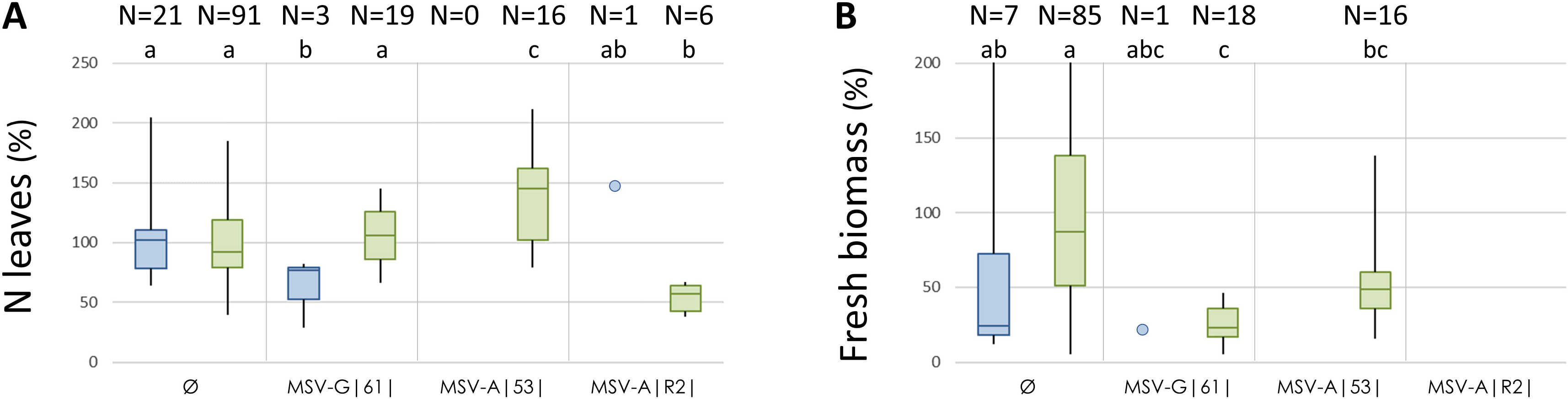
Number of leaves at 28 days post-inoculation (dpi) (**A**) and fresh biomass at 60 dpi (**B**) of MSV infected plants (inoculated with pCAMBIA0380::MSV-G|61|, pCAMBIA0380::MSV-A|53| and pBC-KS::MSV-A|R2|) normalized according to negative control plants (inoculated with pCAMBIA0380::Ø) for *O. sativa indica* cv. IR64 (blue) and *O. glaberrima* cv. Tog5673 (green). Values obtained from 2 independent experiments (A) or only one experiment (B). The numbers of plants used to obtain these data are indicated above. The statistically identical groups are mentioned (a,b,c). The values obtained with the unique IR64 infected after pBC-KS::MSV-A|R2| and pCAMBIA0380::MSV-G|61| agroinoculations are reported by blue points.

## TABLES

**Table Supp.1:** List of wild plant species identified in the borders of 6 rice fields from 2017. The frequency of each plant species was estimated (0: absent; 0.5: rare; 1: covering area < 10%; 2: covering area between 10% and 20%; 3: covering area between 20% and 30%). Samples from the five more frequent plant species have been collected for each field for further analyses.

| Family | Species | 17BM02 | 17BL02 | 17BZ11 | 17SZ07 | 17KA01 | 17TG01 |
| --- | --- | --- | --- | --- | --- | --- | --- |
| Acanthaceae | *Nelsonia canescens* (Lam.) Spreng. | 0.5 | 0 | 0.5 | 1 | 0.5 | 0 |
| Adiantaceae | *Ceratopteris cornuta* (Bauv.) Lepr. | 0 | 1 | 0 | 0 | 0 | 0 |
| Amaranthaceae | *Alternanthera sessilis* (Lin.) R. BR.. | 0 | 0 | 0.5 | 0 | 0 | 0 |
| Amaranthaceae | *Amaranthus graecizans* Lin. | 0 | 0 | 0.5 | 0 | 0 | 0 |
| Amaranthaceae | *Gomphrena celosioides* Mart | 0 | 0 | 0.5 | 2 | 0 | 0 |
| Asteraceae | *Acmeilla caulirhiza* Delile | 0.5 | 0.5 | 0 | 0 | 0.5 | 0 |
| Asteraceae | *Ageratum conyzoides* Lin. | 1 | 0 | 0 | 1 | 0.5 | 0 |
| Asteraceae | *Tridax procumbens* Lin. | 0.5 | 0 | 1 | 0 | 0.5 | 0 |
| Boraginaceae | *Heliotropium indicum* Dc. | 0 | 0 | 0.5 | 0 | 0 | 0 |
| Caesalpiniaceae | *Cassia mimosoides* Lin. | 0 | 0 | 0 | 0.5 | 0 | 0 |
| Caesalpiniaceae | *Cassia tora* Lin. | 0 | 0 | 0.5 | 0.5 | 0 | 0 |
| Campanulaceae | *Sphenoclea zeylanica* Gaertn. | 0.5 | 0.5 | 0 | 0 | 0 | 0 |
| Caparidaceae | *Cleome rutidosperma* DC. | 0.5 | 0 | 0 | 0 | 0 | 0 |
| Caparidaceae | *Cleome viscosa* Lin. | 0 | 0 | 0.5 | 0 | 0 | 0 |
| *Commelinaceae* | *Aneilema umbrosum* Kunth. | 0 | 0 | 0 | 0 | 0 | 0.5 |
| *Commelinaceae* | *Commelina benghalensis* Lin. | 0 | 0 | 0 | 0.5 | 0.5 | 0 |
| *Commelinaceae* | *Commelina forskalaei* Vahl. | 0.5 | 0 | 0 | 0.5 | 0 | 0 |
| *Convolvulaceae* | *Ipomoea aquatica* Forsk. | 0.5 | 0 | 0 | 0 | 0.5 | 0 |
| *Convolvulaceae* | *Ipomoea eriocarpa* R. Br. | 0.5 | 0 | 0.5 | 0.5 | 0 | 0.5 |
| *Cyperaceae* | *Cyperus haspan* Lin. | 0 | 2 | 0 | 0 | 0 | 0 |
| *Cyperaceae* | *Cyperus iria* Lin. | 0 | 0.5 | 0 | 0.5 | 0 | 0 |
| *Cyperaceae* | *Cyperus rotundus* Lin. | 0 | 0 | 0 | 0 | 0.5 | 0 |
| *Cyperaceae* | *Fimbristylis dichotoma* (Lin.) Vahl. | 0.5 | 3 | 0.5 | 2 | 0.5 | 0 |
| *Cyperaceae* | *Kyllinga erecta* Schum. Et Thonn. | 0 | 0 | 0 | 0 | 0.5 | 0 |
| *Cyperaceae* | *Scleria naumanniana* Böck. | 0 | 1 | 0 | 0 | 0 | 0 |
| *Euphorbiaceae* | *Euphorbia herterophylla* Lin. | 0 | 0 | 0 | 0 | 0.5 | 0 |
| *Euphorbiaceae* | *Euphorbia hirta* Lin. | 0.5 | 0 | 0.5 | 0.5 | 0.5 | 0 |
| *Euphorbiaceae* | *Euphorbia prostrata* Ait. | 0 | 0 | 0.5 | 0 | 0 | 0 |
| *Euphorbiaceae* | *Phyllanthus amarus* Schum. Et Thonn. | 0.5 | 0 | 0 | 0 | 0.5 | 0 |
| *Fabaceae* | *Aeschynomene sensitiva* Sw. | 2 | 0.5 | 0 | 2 | 0.5 | 0.5 |
| *Fabaceae* | *Alysicarpus ovalifolius* (S. & Th.)Léon | 0 | 0 | 0 | 0.5 | 0 | 0 |
| *Fabaceae* | *Crotalaria retusa* Lin. | 0 | 0 | 0 | 0.5 | 0 | 0 |
| *Fabaceae* | *Desmodium incanum* | 3 | 0 | 0 | 0 | 0 | 0 |
| *Fabaceae* | *Desmodium salicifolium* (Poir) D C. | 0 | 0 | 0 | 0.5 | 0.5 | 0 |
| *Fabaceae* | *Desmodium triflorum* (Lin.) DC | 3 | 0 | 0 | 0 | 0 | 0 |
| *Fabaceae* | *Sesbania sesban* (Lin.) Merrill | 0 | 0.5 | 0 | 0 | 0 | 0 |
| *Lamiaceae* | *Hyptis specigera* Lam. | 0 | 0.5 | 0 | 2 | 0 | 0.5 |
| *Lamiaceae* | *Ocimum Americanum* Lin. | 0 | 0 | 0.5 | 0.5 | 0 | 0.5 |
| *Malvaceae* | *Sida acuta* Burm. F. | 0.5 | 0 | 0 | 0.5 | 0 | 0 |
| *Malvaceae* | *Sida alba* Lin. | 0.5 | 0 | 0 | 0 | 0 | 0 |
| *Malvaceae* | *Sida ovata* Forsk. | 1 | 0 | 0 | 0.5 | 0 | 0 |
| *Malvaceae* | *Sida urens* Lin. | 0 | 0 | 0.5 | 0.5 | 0 | 0 |
| *Nyctaginaceae* | *Boehraavia vulvarifolia* Poir | 0 | 0 | 1 | 0 | 0 | 0 |
| *Oenotheraceae* | *Ludwigia decurrens* Wal. | 0.5 | 1 | 0 | 0.5 | 0 | 0 |
| *Oenotheraceae* | *Ludwigia erecta* | 0.5 | 1 | 0 | 1 | 0.5 | 0 |
| *Passifloraceae* | *Passiflora foetida* Lin. | 0 | 0 | 0 | 0 | 1 | 0 |
| *Pedaliaceae* | *Ceratotheca sesamoides* Endl. | 0 | 0 | 0 | 0 | 0 | 0.5 |
| *Poaceae* | *Andropogon pseudapricus* sStapf. | 0 | 0 | 0 | 0 | 0 | 0.5 |
| *Poaceae* | *Brachiaria lata* (Chum.) Hubb. | 0 | 0 | 0 | 0 | 0 | 0.5 |
| *Poaceae* | *Brachiaria xantholeuca* Stapf | 0 | 0 | 0 | 0 | 0.5 | 0 |
| *Poaceae* | *Dactyloctenium aegyptium* Beauv. | 0 | 0 | 0 | 1 | 0 | 0.5 |
| *Poaceae* | *Digitaria horizontalis* | 0 | 0 | 0 | 0 | 0 | 0.5 |
| *Poaceae* | *Echinochloa colona* Link. | 0 | 0 | 0.5 | 0.5 | 0.5 | 0 |
| *Poaceae* | *Echinochloa stagnina* P. Beauv. | 1 | 0.5 | 0 | 0 | 0 | 0 |
| *Poaceae* | *Eichhornia crassipes* (Mart.) Solms. | 0 | 0.5 | 0 | 0 | 0 | 0 |
| *Poaceae* | *Eleusine indica* Gaertn. | 0 | 0.5 | 1 | 0 | 0.5 | 0 |
| *Poaceae* | *Eragrostis gangetica* Steud. | 0 | 2 | 0 | 0 | 0 | 0 |
| *Poaceae* | *Eragrostis tremula* Hochst. | 0 | 0 | 0 | 0 | 0 | 0.5 |
| *Poaceae* | *Imperata cylindrica* (Lin.) Raeusch. | 0 | 0 | 1 | 0 | 0.5 | 0 |
| *Poaceae* | *Leersia hexandra* | 0.5 | 0 | 0 | 0 | 0 | 0 |
| Poaceae | *Oryza barthii* A. Chev. | 0 | 3 | 0 | 0.5 | 0 | 0 |
| Poaceae | *Paspalum scrobiculatum* Lin. | 0.5 | 2 | 0.5 | 3 | 0.5 | 0 |
| Poaceae | *Pennisetum pedicellatum* Trin. | 0 | 0 | 0 | 0.5 | 0.5 | 0.5 |
| Poaceae | *Rottboellia exaltata* Lin. | 0.5 | 0 | 0 | 2 | 0 | 0 |
| Poaceae | *Sacciolepis africana* Hubb. Et Snowden | 0 | 2 | 0 | 0 | 0 | 0 |
| Poaceae | *Schizachyrium brevifolium* Nees. | 0 | 1 | 0 | 3 | 0 | 0 |
| Poaceae | *Setaria palledifusca* Stapf et Hubb. | 0 | 0 | 0 | 0 | 0.5 | 0.5 |
| Portulacaceae | *Portulaca oleracea* Lin | 0 | 0 | 0.5 | 0 | 0 | 0 |
| Rubiaceae | *Mitracarpus scaber* Zucc. | 0 | 0 | 0 | 0 | 0 | 0.5 |
| Rubiaceae | *Oldenlandia herbacea* (Lin.) Roxb. | 0 | 0 | 0 | 0.5 | 0 | 0.5 |
| Rubiaceae | *Spermacoce stachydea* DC. | 0 | 0 | 0.5 | 0 | 0 | 0 |
| Rubiaceae | *Spermacoce vertillata* Lin. | 3 | 0 | 0 | 2 | 0.5 | 0 |
| Scrophulariaceae | *Rhamphicarpa fistulosa* (Hochst.) Benth. | 0 | 0.5 | 0 | 0 | 0 | 0 |
| Solanaceae | *Physalis angulata* Lin. | 0 | 0 | 0 | 0 | 0.5 | 0 |
| Solanaceae | *Schwenckia americana* Lin. | 0 | 0 | 0 | 0 | 0 | 0 |
| Sterculiaceae | *Melochia cordifolia* Lin. | 0 | 1 | 0 | 0.5 | 0 | 0 |
| Tiliaceae | *Corchorus olitorius* Lin. | 0 | 0 | 0 | 0.5 | 0 | 0 |
| Tiliaceae | *Triumfetta rhomboidea* Jacq | 0 | 0 | 0 | 0.5 | 0 | 0 |
| Verbenaceae | *Stachytarpheta angustifolia* (Mill.) Vahl. | 0 | 0 | 0 | 0 | 0.5 | 0 |

**Table Supp.2:**
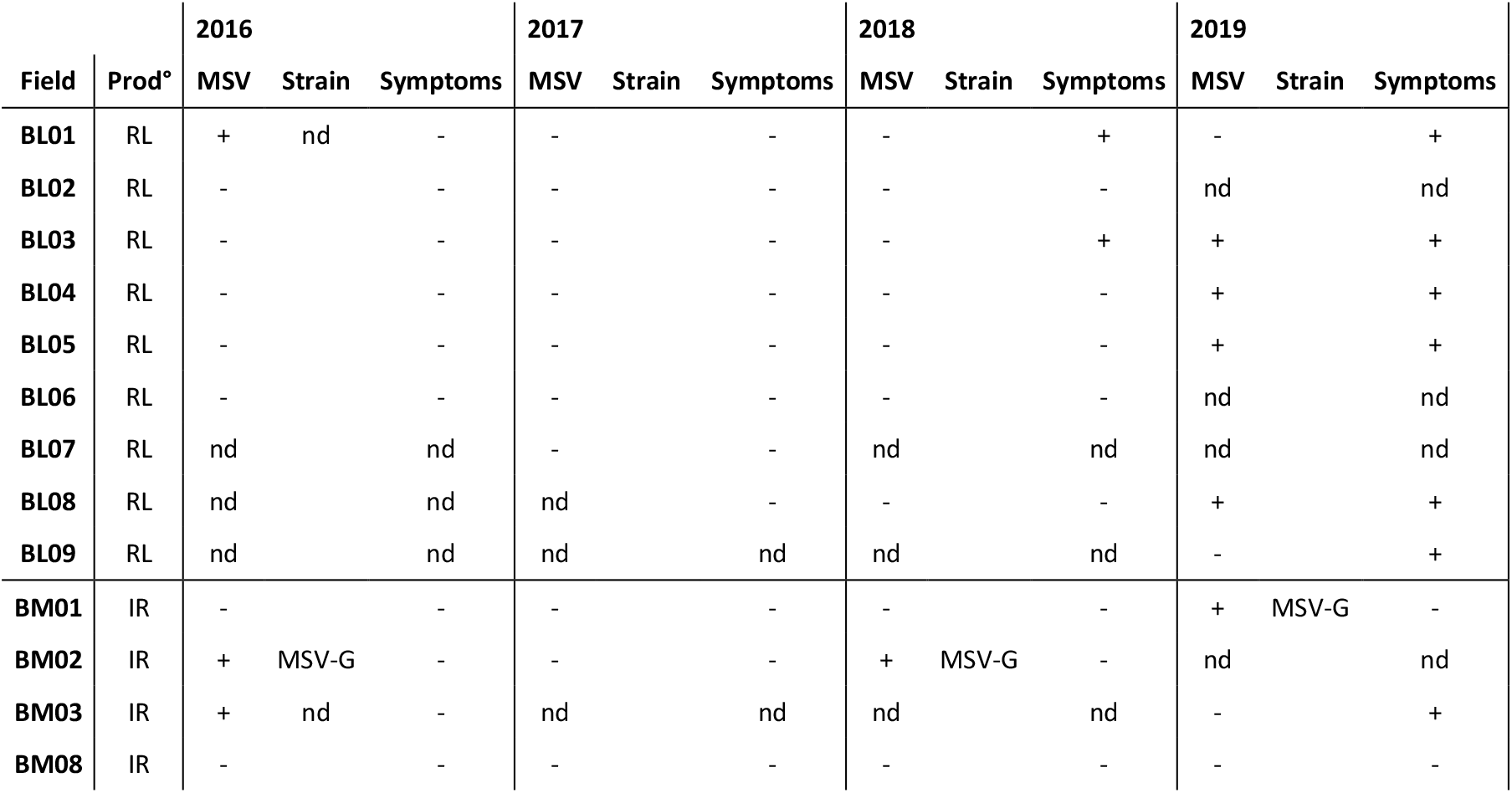
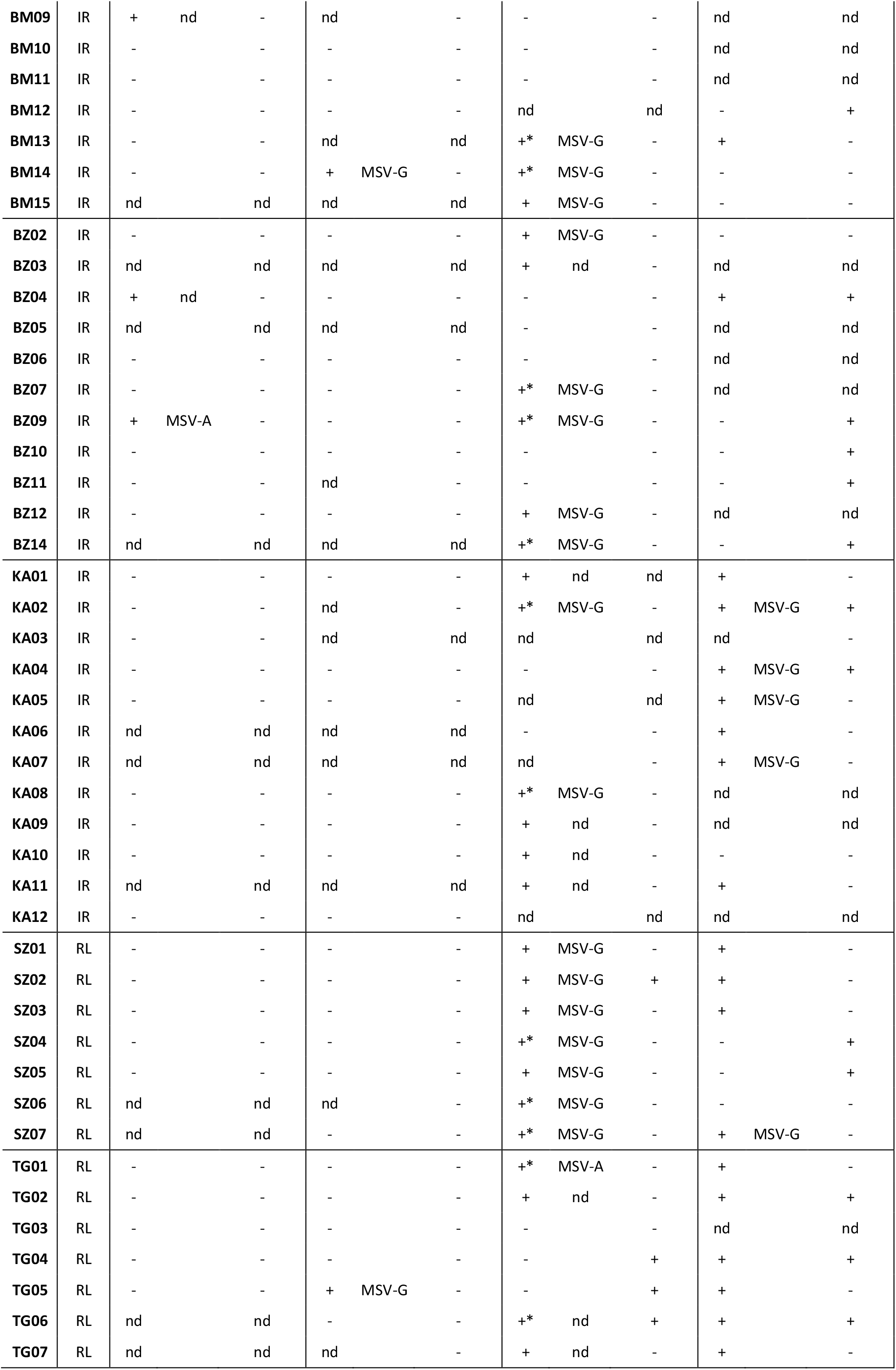
Detection of MSV in rice fields from 6 sites (BM: Bama; BZ: Banzon; KA: Karfiguela; BL: Badala; SZ: Senzon; TG: Tengrela) representing two rice production systems (IR: irrigated; RL: rainfed lowland) in 2016, 2017, 2018 and 2019. The MSV strain (MSV-G or MSV-A) is indicated when identified, and the observation of presence (+) or absence (-) of symptoms putatively related to MSV infection in each rice field is reported. The fields for which the MSV prevalence has been estimated are indicated (*). nd: not determined.

**Table Supp.3:**
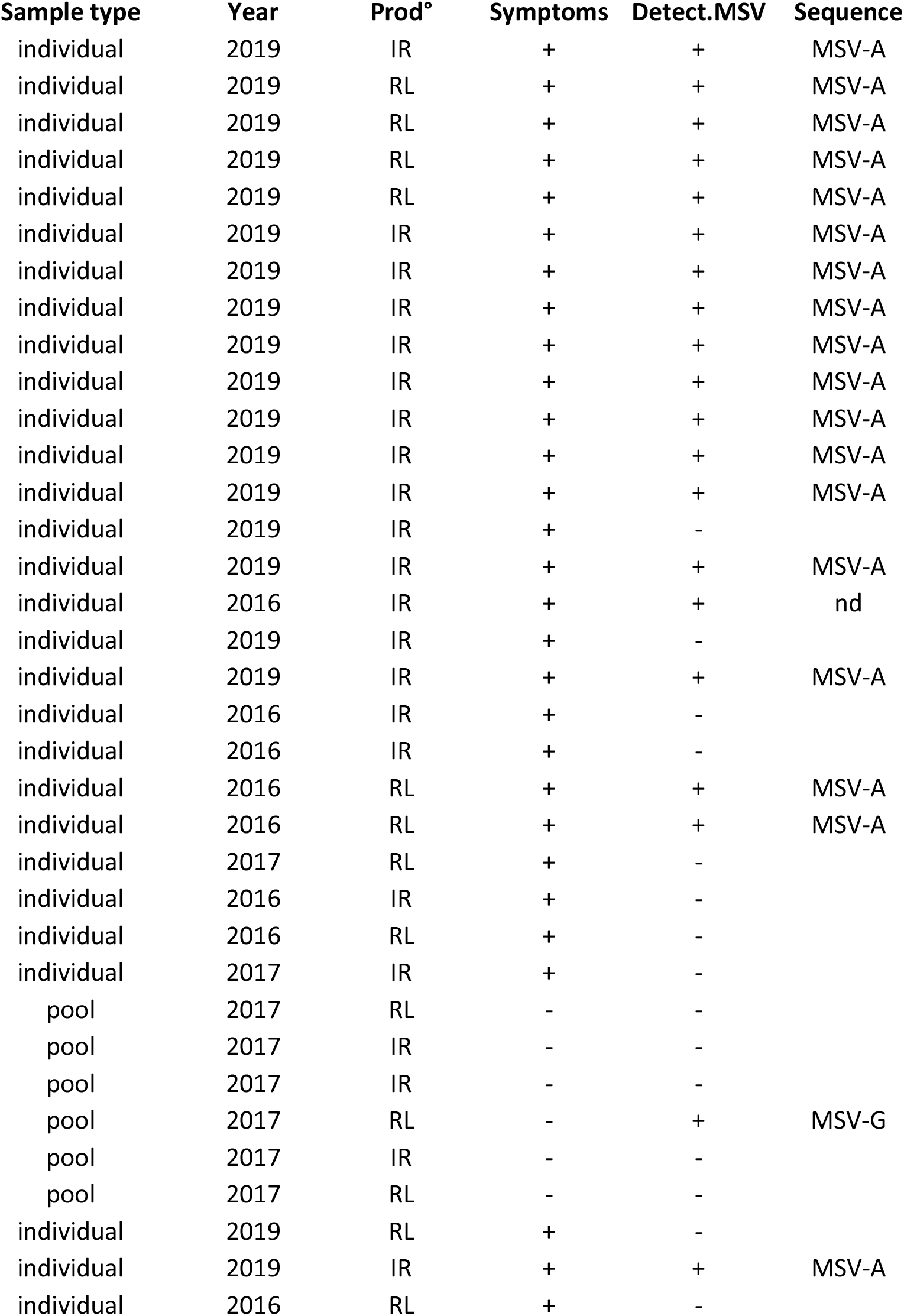
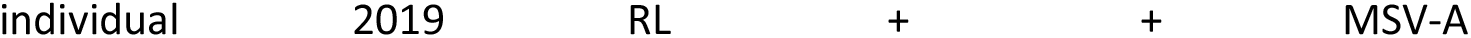
Samples of cultivated (maize, sugarcane) and wild grasses analyzed during this study. The sample type (individual: from an individual plant; pool: mixed leaf samples from several plants), the rice production systems where these plants were collected (IR: irrigated; RL: rainfed lowland), the presence (+) or absence (-) of symptoms putatively related to MSV infection, the results of PCR detection of MSV and the MSV strain (MSV-G or MSV-A) identified for each sample are indicated. nd: not determined.

**Table Supp.4:** Isolate names, origins, sequencing methods and accession numbers of complete MSV genomes obtained during this study.

| Sample name | Host species | Compartment | Year | Sample type | MSV | Seq. Method | Accession n° |
| --- | --- | --- | --- | --- | --- | --- | --- |
| 16BZ09\|53\| | *Oryza sativa* | cultivated | 2016 | pool | MSV-A | Sanger | OR258397 |
| 17BM11\|m61\| | *Oryza sativa* | cultivated | 2017 | pool | MSV-G | Cloning, Sanger | OR258400 |
| 17BM11\|Contig1671\| | *Oryza sativa* | cultivated | 2017 | pool | MSV-G | VANA-Illumina | - |
| 17TG05\|Contig844\| | *Oryza sativa* | cultivated | 2017 | pool | MSV-G | VANA-Illumina | OR258402 |
| 18TG01\|110\| | *Oryza sativa* | cultivated | 2018 | pool | MSV-A | Sanger | OR258399 |
| 18BZ14\|102\| | *Oryza sativa* | cultivated | 2018 | pool | MSV-G | Sanger | OR258398 |
| 19BL\|5\|Maize | *Zea mays* | cultivated | 2019 | individual | MSV-A | Sanger | OR258387 |
| 19BL\|7\|Maize | *Zea mays* | cultivated | 2019 | individual | MSV-A | Sanger | OR258388 |
| 19KA\|12\|Sugarcane | *Saccharum sp.* | cultivated | 2019 | individual | MSV-A | Sanger | OR258390 |
| 19KA\|13\|Sugarcane | *Saccharum sp.* | cultivated | 2019 | individual | MSV-A | Sanger | OR258391 |
| 19KA\|16\|Sugarcane | *Saccharum sp.* | cultivated | 2019 | individual | MSV-A | Sanger | OR258393 |
| 19KA\|17\|Sugarcane | *Saccharum sp.* | cultivated | 2019 | individual | MSV-A | Sanger | OR258394 |
| 16SZ\|22\|Eragrostis | *Eragrostis sp.* | wild | 2016 | individual | MSV-A | Sanger | OR258396 |
| 19BM\|2\|Echinochloa | *Echinochloa colona* | wild | 2019 | individual | MSV-A | Sanger | OR258386 |
| 16SZ\|21\|O.longistaminata | *Oryza longistaminata* | wild | 2016 | individual | MSV-A | Sanger | OR258395 |
| 19KA\|8\|Poaceae | *Poaceae ?* | wild | 2019 | individual | MSV-A | Sanger | OR258389 |
| 19SZ\|14\|Poaceae | *Poaceae ?* | wild | 2019 | individual | MSV-A | Sanger | OR258392 |
| 17TG01\|m96\| | *Poaceae ?* | wild | 2017 | pool | MSV-G | Sanger | OR258401 |
| 17TG01\|Contig711\| | *Poaceae ?* | wild | 2017 | pool | MSV-G | VANA-Illumina | - |

**Table Supp.5:** Detection of MSV (MSV-G or MSV-A) in the 15-16 individual plants collected without *a priori* over a grid in 2018 in 12 rice fields. The position of each sample over the grid (cf. Figure Supp1), the rice production systems (IR: irrigated; RL: rainfed lowland), the results of MSV detection by PCR and the symptom putatively related to MSV infection and observed for each plant in field are indicated.

| n° plant | Field | Position | Prod° | Detect.MSV | Symptoms |
| --- | --- | --- | --- | --- | --- |
| MP2808 | 18-BM13 | A1 | IR | - | - |
| MP2809 | 18-BM13 | A2 | IR | + (MSV - G ) | + |
| MP2810 | 18-BM13 | A3 | IR | + (MSV - G ) | + |
| MP2811 | 18-BM13 | A4 | IR | - | + |
| MP2812 | 18-BM13 | B1 | IR | - | - |
| MP2813 | 18-BM13 | B2 | IR | - | - |
| MP2814 | 18-BM13 | B3 | IR | - | - |
| MP2815 | 18-BM13 | B4 | IR | - | - |
| MP2816 | 18-BM13 | C1 | IR | + (MSV - G ) | - |
| MP2817 | 18-BM13 | C2 | IR | - | - |
| MP2818 | 18-BM13 | C3 | IR | + | - |
| MP2819 | 18-BM13 | C4 | IR | - | - |
| MP2820 | 18-BM13 | D1 | IR | + (MSV - G ) | - |
| MP2821 | 18-BM13 | D2 | IR | - | + |
| MP2822 | 18-BM13 | D3 | IR | - | - |
| MP2823 | 18-BM13 | D4 | IR | + | + |
| MP2824 | 18-BM14 | A1 | IR | - | - |
| MP2825 | 18-BM14 | A2 | IR | - | - |
| MP2826 | 18-BM14 | A3 | IR | + | - |
| MP2827 | 18-BM14 | A4 | IR | + | - |
| MP2828 | 18-BM14 | B1 | IR | + | - |
| MP2829 | 18-BM14 | B2 | IR | - | - |
| MP2830 | 18-BM14 | B3 | IR | + | - |
| MP2831 | 18-BM14 | B4 | IR | - | - |
| MP2832 | 18-BM14 | C1 | IR | + | - |
| MP2833 | 18-BM14 | C2 | IR | - | - |
| MP2834 | 18-BM14 | C3 | IR | - | - |
| MP2835 | 18-BM14 | C4 | IR | - | - |
| MP2836 | 18-BM14 | D1 | IR | - | - |
| MP2837 | 18-BM14 | D2 | IR | - | - |
| MP2838 | 18-BM14 | D3 | IR | - | - |
| MP2839 | 18-BM14 | D4 | IR | - | - |
| MP2936 | 18-BZ07 | A1 | IR | - | - |
| MP2937 | 18-BZ07 | A2 | IR | - | - |
| MP2938 | 18-BZ07 | A3 | IR | + | - |
| MP2939 | 18-BZ07 | A4 | IR | + | - |
| MP2940 | 18-BZ07 | B1 | IR | - | - |
| MP2941 | 18-BZ07 | B2 | IR | + | - |
| MP2942 | 18-BZ07 | B3 | IR | + | - |
| MP2943 | 18-BZ07 | B4 | IR | - | - |
| MP2944 | 18-BZ07 | C1 | IR | - | - |
| MP2945 | 18-BZ07 | C2 | IR | - | - |
| MP2946 | 18-BZ07 | C3 | IR | nd | - |
| MP2947 | 18-BZ07 | C4 | IR | - | - |
| MP2948 | 18-BZ07 | D1 | IR | - | - |
| MP2949 | 18-BZ07 | D2 | IR | - | - |
| MP2950 | 18-BZ07 | D3 | IR | - | - |
| MP2951 | 18-BZ07 | D4 | IR | - | - |
| MP2952 | 18-BZ09 | A1 | IR | + | - |
| MP2953 | 18-BZ09 | A2 | IR | + | - |
| MP2954 | 18-BZ09 | A3 | IR | - | - |
| MP2955 | 18-BZ09 | A4 | IR | + | - |
| MP2956 | 18-BZ09 | B1 | IR | - | - |
| MP2957 | 18-BZ09 | B2 | IR | + | - |
| MP2958 | 18-BZ09 | B3 | IR | + | - |
| MP2959 | 18-BZ09 | B4 | IR | + | - |
| MP2960 | 18-BZ09 | C1 | IR | + | - |
| MP2961 | 18-BZ09 | C2 | IR | + | - |
| MP2962 | 18-BZ09 | C3 | IR | - | - |
| MP2963 | 18-BZ09 | C4 | IR | + | - |
| MP2964 | 18-BZ09 | D1 | IR | - | - |
| MP2965 | 18-BZ09 | D2 | IR | - | - |
| MP2966 | 18-BZ09 | D3 | IR | + | - |
| MP2967 | 18-BZ09 | D4 | IR | - | - |
| MP3016 | 18-BZ14 | A1 | IR | + | - |
| MP3017 | 18-BZ14 | A2 | IR | - | - |
| MP3018 | 18-BZ14 | A3 | IR | - | - |
| MP3019 | 18-BZ14 | A4 | IR | - | - |
| MP3020 | 18-BZ14 | B1 | IR | - | - |
| MP3021 | 18-BZ14 | B2 | IR | - | - |
| MP3022 | 18-BZ14 | B3 | IR | - | - |
| MP3023 | 18-BZ14 | B4 | IR | + (MSV - G ) | - |
| MP3024 | 18-BZ14 | C1 | IR | - | - |
| MP3025 | 18-BZ14 | C2 | IR | - | - |
| MP3026 | 18-BZ14 | C3 | IR | - | - |
| MP3027 | 18-BZ14 | C4 | IR | - | - |
| MP3028 | 18-BZ14 | D1 | IR | - | - |
| MP3029 | 18-BZ14 | D2 | IR | - | - |
| MP3030 | 18-BZ14 | D3 | IR | - | - |
| MP3031 | 18-BZ14 | D4 | IR | + (MSV - G ) | - |
| MP3064 | 18-SZ04 | A1 | RL | + | - |
| MP3065 | 18-SZ04 | A2 | RL | - | - |
| MP3066 | 18-SZ04 | A3 | RL | - | - |
| MP3067 | 18-SZ04 | A4 | RL | - | - |
| MP3068 | 18-SZ04 | B1 | RL | + | - |
| MP3069 | 18-SZ04 | B2 | RL | + | - |
| MP3070 | 18-SZ04 | B3 | RL | - | - |
| MP3071 | 18-SZ04 | B4 | RL | + | - |
| MP3072 | 18-SZ04 | C1 | RL | + | - |
| MP3073 | 18-SZ04 | C2 | RL | - | - |
| MP3074 | 18-SZ04 | C3 | RL | - | - |
| MP3075 | 18-SZ04 | C4 | RL | - | - |
| MP3076 | 18-SZ04 | D1 | RL | + | - |
| MP3077 | 18-SZ04 | D2 | RL | - | - |
| MP3078 | 18-SZ04 | D3 | RL | + | - |
| MP3079 | 18-SZ04 | D4 | RL | + | - |
| MP3096 | 18-SZ06 | A1 | RL | - | - |
| MP3097 | 18-SZ06 | A2 | RL | - | - |
| MP3098 | 18-SZ06 | A3 | RL | - | - |
| MP3099 | 18-SZ06 | A4 | RL | + (MSV - G ) | - |
| MP3100 | 18-SZ06 | B1 | RL | - | - |
| MP3101 | 18-SZ06 | B2 | RL | - | - |
| MP3102 | 18-SZ06 | B3 | RL | - | - |
| MP3103 | 18-SZ06 | B4 | RL | - | - |
| MP3104 | 18-SZ06 | C1 | RL | - | - |
| MP3105 | 18-SZ06 | C2 | RL | + (MSV - G ) | - |
| MP3106 | 18-SZ06 | C3 | RL | - | - |
| MP3107 | 18-SZ06 | C4 | RL | - | - |
| MP3108 | 18-SZ06 | D1 | RL | - | - |
| MP3109 | 18-SZ06 | D2 | RL | + | - |
| MP3110 | 18-SZ06 | D3 | RL | + | - |
| MP3111 | 18-SZ06 | D4 | RL | + | - |
| MP3112 | 18-SZ07 | A1 | RL | nd | - |
| MP3113 | 18-SZ07 | A2 | RL | - | - |
| MP3114 | 18-SZ07 | A3 | RL | - | - |
| MP3115 | 18-SZ07 | A4 | RL | - | - |
| MP3116 | 18-SZ07 | B1 | RL | - | - |
| MP3117 | 18-SZ07 | B2 | RL | - | - |
| MP3118 | 18-SZ07 | B3 | RL | + | - |
| MP3119 | 18-SZ07 | B4 | RL | - | - |
| MP3120 | 18-SZ07 | C1 | RL | - | - |
| MP3121 | 18-SZ07 | C2 | RL | + | - |
| MP3122 | 18-SZ07 | C3 | RL | - | - |
| MP3123 | 18-SZ07 | C4 | RL | - | - |
| MP3124 | 18-SZ07 | D1 | RL | + | - |
| MP3125 | 18-SZ07 | D2 | RL | + | - |
| MP3126 | 18-SZ07 | D3 | RL | + | - |
| MP3127 | 18-SZ07 | D4 | RL | - | - |
| MP3160 | 18-KA02 | A1 | IR | - | - |
| MP3161 | 18-KA02 | A2 | IR | + | - |
| MP3162 | 18-KA02 | A3 | IR | + | - |
| MP3163 | 18-KA02 | A4 | IR | - | - |
| MP3164 | 18-KA02 | B1 | IR | - | - |
| MP3165 | 18-KA02 | B2 | IR | - | - |
| MP3166 | 18-KA02 | B3 | IR | + | - |
| MP3167 | 18-KA02 | B4 | IR | - | - |
| MP3168 | 18-KA02 | C1 | IR | - | - |
| MP3169 | 18-KA02 | C2 | IR | - | - |
| MP3170 | 18-KA02 | C3 | IR | - | - |
| MP3171 | 18-KA02 | C4 | IR | - | - |
| MP3172 | 18-KA02 | D1 | IR | - | - |
| MP3173 | 18-KA02 | D2 | IR | - | - |
| MP3174 | 18-KA02 | D3 | IR | - | - |
| MP3175 | 18-KA02 | D4 | IR | + | - |
| MP3224 | 18-KA08 | A1 | IR | - | - |
| MP3225 | 18-KA08 | A2 | IR | + | - |
| MP3226 | 18-KA08 | A3 | IR | + (MSV - G ) | - |
| MP3227 | 18-KA08 | A4 | IR | - | - |
| MP3228 | 18-KA08 | B1 | IR | - | - |
| MP3229 | 18-KA08 | B2 | IR | - | - |
| MP3230 | 18-KA08 | B3 | IR | - | - |
| MP3231 | 18-KA08 | B4 | IR | - | - |
| MP3232 | 18-KA08 | C1 | IR | + (MSV - G ) | - |
| MP3233 | 18-KA08 | C2 | IR | - | - |
| MP3234 | 18-KA08 | C3 | IR | + (MSV - G ) | - |
| MP3235 | 18-KA08 | C4 | IR | - | - |
| MP3236 | 18-KA08 | D1 | IR | + (MSV - G ) | - |
| MP3237 | 18-KA08 | D2 | IR | - | - |
| MP3238 | 18-KA08 | D3 | IR | - | - |
| MP3239 | 18-KA08 | D4 | IR | - | - |
| MP3288 | 18-TG01 | A1 | RL | - | - |
| MP3289 | 18-TG01 | A2 | RL | - | - |
| MP3290 | 18-TG01 | A3 | RL | - | - |
| MP3291 | 18-TG01 | A4 | RL | - | - |
| MP3292 | 18-TG01 | B1 | RL | - | - |
| MP3293 | 18-TG01 | B2 | RL | - | - |
| MP3294 | 18-TG01 | B3 | RL | + | - |
| MP3295 | 18-TG01 | B4 | RL | - | - |
| MP3296 | 18-TG01 | C1 | RL | - | - |
| MP3297 | 18-TG01 | C2 | RL | - | - |
| MP3298 | 18-TG01 | C3 | RL | - | - |
| MP3299 | 18-TG01 | C4 | RL | - | - |
| MP3300 | 18-TG01 | D1 | RL | - | - |
| MP3301 | 18-TG01 | D2 | RL | - | - |
| MP3302 | 18-TG01 | D3 | RL | - | - |
| MP3303 | 18-TG01 | D4 | RL | - | - |
| MP3368 | 18-TG06 | A1 | RL | - | - |
| MP3369 | 18-TG06 | A2 | RL | - | - |
| MP3370 | 18-TG06 | A3 | RL | - | - |
| MP3371 | 18-TG06 | A4 | RL | - | - |
| MP3372 | 18-TG06 | B1 | RL | - | - |
| MP3373 | 18-TG06 | B2 | RL | - | - |
| MP3374 | 18-TG06 | B3 | RL | - | - |
| MP3375 | 18-TG06 | B4 | RL | + | - |
| MP3376 | 18-TG06 | C1 | RL | - | - |
| MP3377 | 18-TG06 | C2 | RL | + | - |
| MP3378 | 18-TG06 | C3 | RL | - | + |
| MP3379 | 18-TG06 | C4 | RL | - | - |
| MP3380 | 18-TG06 | D1 | RL | + | - |
| MP3381 | 18-TG06 | D2 | RL | - | - |
| MP3382 | 18-TG06 | D3 | RL | + | - |
| MP3383 | 18-TG06 | D4 | RL | + | - |

**Table Supp.6:** Number of non-symptomatic (-) and symptomatic (+) plants observed 28 days post-inoculation (dpi) of *Agrobacterium tumefaciens* strains C58C1 and EHA105 transformed with pCAMBIA0380::Ø (negative control), pCAMBIA0380::MSV-G|61|, pCAMBIA0380::MSV-A|53| and pBC-KS::MSV-A|R2| in *Oryza sativa indica* cv. IR64, *O. glaberrima* cv. Tog5673 and *Zea mays* cv. Golden Bantam during two independent experiments. Percentages of symptomatic plants are indicated with %(+). nd: not determined.

| Infectious clones | Agro. | #Exp. | *O. sativa indica*<br>cv. IR64 |  |  | *O. glaberrima* cv.<br>Tog5673 |  |  | *Zea mays* cv. Golden<br>Bantam |  |  |
| --- | --- | --- | --- | --- | --- | --- | --- | --- | --- | --- | --- |
|  |  |  | - | t+ | %(+) | - | + | %(+) | - | + | %(+) |
| pCAMBIA0380::Ø | C58C1 | 1 | 5 | 0 | 0.0 | 4 | 0 | 0.0 | nd | nd | nd |
| pCAMBIA0380::Ø | EHA105 | 1 | 6 | 0 | 0.0 | 5 | 0 | 0.0 | nd | nd | nd |
| pCAMBIA0380::Ø | EHA105 | 2 | 5 | 0 | 0.0 | 11 | 0 | 0.0 | 3 | 0 | 0.0 |
| pCAMBIA0380::MSV-G\|61\| | C58C1 | 1 | 33 | 0 | 0.0 | 30 | 1 | 3.2 | 2 | 0 | 0.0 |
| pCAMBIA0380::MSV-G\|61\| | EHA105 | 1 | 34 | 2 | 5.6 | 36 | 1 | 2.7 | 2 | 0 | 0.0 |
| pCAMBIA0380::MSV-G\|61\| | EHA105 | 2 | 131 | 1 | 0.8 | 119 | 18 | 13.1 | 80 | 0 | 0.0 |
| pCAMBIA0380::MSV-A\|53\| | EHA105 | 2 | 127 | 0 | 0.0 | 110 | 16 | 12.7 | 67 | 1 | 1.5 |
| pBCKS::MSV-A\|R2\| | C58C1 | 1 | 26 | 0 | 0.0 | 27 | 5 | 15.6 | 2 | 1 | 33.3 |
| pBCKS::MSV-A\|R2\| | EHA105 | 1 | 23 | 1 | 4.2 | 28 | 1 | 3.4 | 2 | 0 | 0.0 |
| MSV-G\|61\| |  | 1+2 | 198 | 3 | 1.5 | 185 | 20 | 9.8 | 84 | 0 | 0.0 |
| MSV-A\|53\| |  | 2 | 127 | 0 | 0.0 | 110 | 16 | 12.7 | 67 | 1 | 1.5 |
| MSV-A\|R2\| |  | 1 | 49 | 1 | 2.0 | 55 | 6 | 9.8 | 4 | 1 | 20.0 |
| MSV-G\|61\| |  | 1+2 | 198 | 3 | 1.5 | 185 | 20 | 9.8 | 84 | 0 | 0.0 |
| MSV-A\|53+R2\| |  | 1+2 | 176 | 1 | 0.6 | 165 | 22 | 11.8 | 71 | 2 | 2.7 |
| Total MSV-G/A |  | 1+2 | 374 | 4 | 1.1 | 350 | 42 | 10.7 | 155 | 2 | 1.3 |

## REFERENCES

Abo, M.E., Sy, A.A., 1997. Rice Virus Diseases: Epidemiology and Management Strategies. Journal of Sustainable Agriculture 11, 113–134. 10.1300/J064v11n02_09

Anderson, P.K., Cunningham, A.A., Patel, N.G., Morales, F.J., Epstein, P.R., Daszak, P., 2004. Emerging infectious diseases of plants: pathogen pollution, climate change and agrotechnology drivers. TRENDS in Ecology and Evolution 19, 535–544. 10.1016/j.tree.2004.07.021

Asanzi, C.M., Bosque-Perez, N.A., Nault, L.R., 1995. Movement of Cicadulina storeyi (Homoptera: Cicadellidae) in maize fields and its behaviour in relation to maize growth stage. Insect sci. appl. 16, 39–44. 10.1017/S1742758400018300

Bagayoko, I., Celli, M.G., Romay, G., Poulicard, N., Pinel-Galzi, A., Julian, C., Filloux, D., Roumagnac, P., Sérémé, D., Bragard, C., Hébrard, E., 2021. Genetic Diversity of Rice stripe necrosis virus and New Insights into Evolution of the Genus Benyvirus. Viruses 13, 737. 10.3390/v13050737

Baker, R.E., Mahmud, A.S., Miller, I.F., Rajeev, M., Rasambainarivo, F., Rice, B.L., Takahashi, S., Tatem, A.J., Wagner, C.E., Wang, L.-F., Wesolowski, A., Metcalf, C.J.E., 2022. Infectious disease in an era of global change. Nat Rev Microbiol 20, 193–205. 10.1038/s41579-021-00639-z

Barro, M., Kassankogno, A.I., Wonni, I., Sérémé, D., Somda, I., Kaboré, H.K., Béna, G., Brugidou, C., Tharreau, D., Tollenaere, C., 2021a. Spatiotemporal Survey of Multiple Rice Diseases in Irrigated Areas Compared to Rainfed Lowlands in the Western Burkina Faso. Plant Disease 105, 3889–3899. 10.1094/PDIS-03-21-0579-RE

Barro, M., Konate, K.A., Wonni, I., Kassankogno, A.I., Sabot, F., Albar, L., Somda, I., Béna, G., Ghesquière, A., Kam, H., Sié, M., Cubry, P., Tollenaere, C., 2021b. Assessment of Genetic Diversity of Rice in Registered Cultivars and Farmers’ Fields in Burkina Faso. Crops 1, 129–140. 10.3390/crops1030013

Barro, M., Wonni, I., Simonin, M., Kassankogno, A.I., Klonowska, A., Moulin, L., Béna, G., Somda, I., Brunel, C., Tollenaere, C., 2022. The impact of the rice production system (irrigated vs lowland) on root-associated microbiome from farmer’s fields in western Burkina Faso. FEMS Microbiology Ecology 98, fiac085. 10.1093/femsec/fiac085

Bernardo, P., Charles-Dominique, T., Barakat, M., Ortet, P., Fernandez, E., Filloux, D., Hartnady, P., Rebelo, T.A., Cousins, S.R., Mesleard, F., Cohez, D., Yavercovski, N., Varsani, A., Harkins, G.W., Peterschmitt, M., Malmstrom, C.M., Martin, D.P., Roumagnac, P., 2018. Geometagenomics illuminates the impact of agriculture on the distribution and prevalence of plant viruses at the ecosystem scale. ISME J 12, 173–184. 10.1038/ismej.2017.155

Billard, E., Barro, M., Sérémé, D., Bangratz, M., Wonni, I., Koala, M., Kassankogno, A.I., Hébrard, E., Thébaud, G., Brugidou, C., Poulicard, N., Tollenaere, C., 2023. Dynamics of the rice yellow mottle disease in western Burkina Faso: epidemic monitoring, spatio-temporal variation of viral diversity and pathogenicity in a disease hotspot (preprint). Evolutionary Biology. 10.1101/2023.03.27.534376

Boulton, M.I., Buchholz, W.G., Marks, M.S., Markham, P.G., Davies, J.W., 1989. Specificity of Agrobacterium-mediated delivery of maize streak virus DNA to members of the Gramineae. Plant Mol Biol 12, 31–40. 10.1007/BF00017445

Claverie, S., Ouattara, A., Hoareau, M., Filloux, D., Varsani, A., Roumagnac, P., Martin, D.P., Lett, J.-M., Lefeuvre, P., 2019. Exploring the diversity of Poaceae-infecting mastreviruses on Reunion Island using a viral metagenomics-based approach. Sci Rep 9, 12716. 10.1038/s41598-019-49134-9

Cubry, P., Tranchant-Dubreuil, C., Thuillet, A.-C., Monat, C., Ndjiondjop, M.-N., Labadie, K., Cruaud, C., Engelen, S., Scarcelli, N., Rhoné, B., Burgarella, C., Dupuy, C., Larmande, P., Wincker, P., François, O., Sabot, F., Vigouroux, Y., 2018. The Rise and Fall of African Rice Cultivation Revealed by Analysis of 246 New Genomes. Current Biology 28, 2274–2282.e6. 10.1016/j.cub.2018.05.066

Damsteegt, V.D., 1983. Maize Streak Virus: I. Host Range and Vulnerability of Maize Germ Plasm. Plant Dis. 67, 734. 10.1094/PD-67-734

Demont, M., 2013. Reversing urban bias in African rice markets: A review of 19 National Rice Development Strategies. Global Food Security 2, 172–181. 10.1016/j.gfs.2013.07.001

Edgar, R.C., 2004. MUSCLE: multiple sequence alignment with high accuracy and high throughput. Nucleic Acids Research 32, 1792–1797. 10.1093/nar/gkh340

Edgar, R.C., Taylor, J., Lin, V., Altman, T., Barbera, P., Meleshko, D., Lohr, D., Novakovsky, G., Buchfink, B., Al-Shayeb, B., Banfield, J.F., de la Peña, M., Korobeynikov, A., Chikhi, R., Babaian, A., 2022. Petabase-scale sequence alignment catalyses viral discovery. Nature 602, 142–147. 10.1038/s41586-021-04332-2

Excoffier, L., Laval, G., Schneider, S., 2005. Arlequin (version 3.0): integrated software package population genetic. Evolutionary Bioinformatics 1, 47–50.

Fiallo-Olivé, E., Lett, J.-M., Martin, D.P., Roumagnac, P., Varsani, A., Zerbini, F.M., Navas-Castillo, J., 2021. ICTV Virus Taxonomy Profile: Geminiviridae 2021: This article is part of the ICTV Virus Taxonomy Profiles collection. Journal of General Virology 102. 10.1099/jgv.0.001696

François, S., Filloux, D., Frayssinet, M., Roumagnac, P., Martin, D.P., Ogliastro, M., Froissart, R., 2018. Increase in taxonomic assignment efficiency of viral reads in metagenomic studies. Virus Research 244, 230–234. 10.1016/j.virusres.2017.11.011

Fuller, C., 1901. Mealie variegation In: First Report of the Government Entomologist, Natal, 1899–1900. Pietermaritzburg, Natal, South Africa: P. Davis & Sons, Government Printers. 17–19.

Gnacadja, C., Berthouly-Salazar, C., Sall, S.N., Zekraoui, L., Sabot, F., Pegalepo, E., Baboucarr, M., Vieira-Dalode, G., Moreira, J., Soumanou, M.M., Azokpota, P., Sie, M., 2018. Phenotypic and genetic characterization of African rice (Oryza glaberrima Steud). IJAR 6, 1389–1398. 10.21474/IJAR01/6569

Gong, P., Tan, H., Zhao, S., Li, H., Liu, H., Ma, Y., Zhang, X., Rong, J., Fu, X., Lozano-Durán, R., Li, F., Zhou, X., 2021. Geminiviruses encode additional small proteins with specific subcellular localizations and virulence function. Nat Commun 12, 4278. 10.1038/s41467-021-24617-4

Gouy, M., Guindon, S., Gascuel, O., 2010. SeaView Version 4: A Multiplatform Graphical User Interface for Sequence Alignment and Phylogenetic Tree Building. Molecular Biology and Evolution 27, 221–224. 10.1093/molbev/msp259

Greninger, A.L., 2018. A decade of RNA virus metagenomics is (not) enough. Virus Research 244, 218–229. 10.1016/j.virusres.2017.10.014

Grimsley, N., Hohn, T., Davies, J.W., Hohn, B., 1987. Agrobacterium-mediated delivery of infectious maize streak virus into maize plants. Nature 325, 177–179. 10.1038/325177a0

Harkins, G.W., Martin, D.P., Duffy, S., Monjane, A.L., Shepherd, D.N., Windram, O.P., Owor, B.E., Donaldson, L., van Antwerpen, T., Sayed, R.A., Flett, B., Ramusi, M., Rybicki, E.P., Peterschmitt, M., Varsani, A., 2009. Dating the origins of the maize-adapted strain of maize streak virus, MSV-A. Journal of General Virology 90, 3066–3074. 10.1099/vir.0.015537-0

Hébrard, E., Poulicard, N., Rakotomalala, M., 2021. Rice Yellow Mottle Virus (Solemoviridae), in: Encyclopedia of Virology. Elsevier, pp. 675–680. 10.1016/B978-0-12-809633-8.21244-2

Heyraud, F., Matzeit, V., Schaefer, S., Schell, J., Gronenborn, B., 1993. The conserved nonanucleotide motif of the geminivirus stem-loop sequence promotes replicational release of virus molecules from redundant copies. Biochimie 75, 605–615. 10.1016/0300-9084(93)90067-3

Isnard, M., Granier, M., Frutos, R., Reynaud, B., Peterschmitt, M., 1998. Quasispecies nature of three maize streak virus isolates obtained through different modes of selection from a population used to assess response to infection of maize cultivars. Journal of General Virology 79, 3091– 3099. 10.1099/0022-1317-79-12-3091

Jones, R.A.C., 2021. Global Plant Virus Disease Pandemics and Epidemics. Plants 10, 233. 10.3390/plants10020233

Kaboré, K.H., Kassankogno, A.I., Adreit, H., Milazzo, J., Guillou, S., Blondin, L., Chopin, L., Ravel, S., Charriat, F., Barro, M., Tollenaere, C., Lebrun, M.-H., Tharreau, D., 2022. Genetic diversity and structure of Bipolaris oryzae and Exserohilum rostratum populations causing brown spot of rice in Burkina Faso based on genotyping-by-sequencing. Front. Plant Sci. 13, 1022348. 10.3389/fpls.2022.1022348

Konaté, G., Traoré, O., 1992. Les hôtes réservoirs du virus de la striure du maïs (MSV) en zone soudano-sahélienne: identification et distribution spatio-temporelle. Phyto 73, 111–117. 10.7202/706027ar

Koonin, E.V., Dolja, V.V., 2018. Metaviromics: a tectonic shift in understanding virus evolution. Virus Research 246, A1–A3. 10.1016/j.virusres.2018.02.001

Kraberger, S., Saumtally, S., Pande, D., Khoodoo, M.H.R., Dhayan, S., Dookun-Saumtally, A., Shepherd, D.N., Hartnady, P., Atkinson, R., Lakay, F.M., Hanson, B., Redhi, D., Monjane, A.L., Windram, O.P., Walters, M., Oluwafemi, S., Michel-Lett, J., Lefeuvre, P., Martin, D.P., Varsani, A., 2017. Molecular diversity, geographic distribution and host range of monocot-infecting mastreviruses in Africa and surrounding islands. Virus Research 238, 171–178. 10.1016/j.virusres.2017.07.001

Kumar, S., Stecher, G., Li, M., Knyaz, C., Tamura, K., 2018. MEGA X: Molecular Evolutionary Genetics Analysis across Computing Platforms. Molecular Biology and Evolution 35, 1547–1549. 10.1093/molbev/msy096

Lefeuvre, P., Martin, D.P., Elena, S.F., Shepherd, D.N., Roumagnac, P., Varsani, A., 2019. Evolution and ecology of plant viruses. Nat Rev Microbiol 17, 632–644. 10.1038/s41579-019-0232-3

Martin, D.P., Murrell, B., Golden, M., Khoosal, A., Muhire, B., 2015. RDP4: Detection and analysis of recombination patterns in virus genomes. Virus Evolution 1. 10.1093/ve/vev003

Martin, D.P., Shepherd, D.N., 2009. The epidemiology, economic impact and control of maize streak disease. Food Sec. 1, 305–315. 10.1007/s12571-009-0023-1

Monjane, A.L., Dellicour, S., Hartnady, P., Oyeniran, K.A., Owor, B.E., Bezuidenhout, M., Linderme, D., Syed, R.A., Donaldson, L., Murray, S., Rybicki, E.P., Kvarnheden, A., Yazdkhasti, E., Lefeuvre, P., Froissart, R., Roumagnac, P., Shepherd, D.N., Harkins, G.W., Suchard, M.A., Lemey, P., Varsani, A., Martin, D.P., 2020. Symptom evolution following the emergence of maize streak virus. eLife 9, e51984. 10.7554/eLife.51984

Monjane, A.L., Harkins, G.W., Martin, D.P., Lemey, P., Lefeuvre, P., Shepherd, D.N., Oluwafemi, S., Simuyandi, M., Zinga, I., Komba, E.K., Lakoutene, D.P., Mandakombo, N., Mboukoulida, J., Semballa, S., Tagne, A., Tiendrebeogo, F., Erdmann, J.B., van Antwerpen, T., Owor, B.E., Flett, B., Ramusi, M., Windram, O.P., Syed, R., Lett, J.-M., Briddon, R.W., Markham, P.G., Rybicki, E.P., Varsani, A., 2011. Reconstructing the History of Maize Streak Virus Strain A Dispersal To Reveal Diversification Hot Spots and Its Origin in Southern Africa. Journal of Virology 85, 9623–9636. 10.1128/JVI.00640-11

Moubset, O., François, S., Maclot, F., Palanga, E., Julian, C., Claude, L., Fernandez, E., Rott, P., Daugrois, J.-H., Antoine-Lorquin, A., Bernardo, P., Blouin, A.G., Temple, C., Kraberger, S., Fontenele, R.S., Harkins, G.W., Ma, Y., Marais, A., Candresse, T., Chéhida, S.B., Lefeuvre, P., Lett, J.-M., Varsani, A., Massart, S., Ogliastro, M., Martin, D.P., Filloux, D., Roumagnac, P., 2022. Virion-Associated Nucleic Acid-Based Metagenomics: A Decade of Advances in Molecular Characterization of Plant Viruses. Phytopathology® 112, 2253–2272. 10.1094/PHYTO-03-22-0096-RVW

Muhire, B.M., Varsani, A., Martin, D.P., 2014. SDT: A Virus Classification Tool Based on Pairwise Sequence Alignment and Identity Calculation. PLoS ONE 9, e108277. 10.1371/journal.pone.0108277

Oyeniran, K.A., Hartnady, P., Claverie, S., Lefeuvre, P., Monjane, A.L., Donaldson, L., Lett, J.-M., Varsani, A., Martin, D.P., 2021. How virulent are emerging maize-infecting mastreviruses? Arch Virol 166, 955–959. 10.1007/s00705-020-04906-x

Peterschmitt, M., Granier, M., Frutos, R., Reynaud, B., 1996. Infectivity and complete nucleotide sequence of the genome of a genetically distinct strain of maize streak virus from Reunion Island. Archives of Virology 141, 1637–1650. 10.1007/BF01718288

Portères, R., 1970. Primary cradles of agriculture in the African continent. Papers in African Prehistory, Cambridge University Press Fage, J&Olivier, R editions, 43–58.

Reynaud, B., Delatte, H., Peterschmitt, M., Fargette, D., 2009. Effects of temperature increase on the epidemiology of three major vector-borne viruses. Eur J Plant Pathol 123, 269–280. 10.1007/s10658-008-9363-5

Rybicki, E.P., 2015. A Top Ten list for economically important plant viruses. Arch Virol 160, 17–20. 10.1007/s00705-014-2295-9

Rybicki, E.P., Pietersen, G., 1999. Plant Virus Disease Problems in The Developing World, in: Advances in Virus Research. Elsevier, pp. 127–175. 10.1016/S0065-3527(08)60346-2

Savary, S., Willocquet, L., Pethybridge, S.J., Esker, P., McRoberts, N., Nelson, A., 2019. The global burden of pathogens and pests on major food crops. Nat Ecol Evol 3, 430–439. 10.1038/s41559-018-0793-y

Sereme, D., Neya, B.J., Bangratz, M., Brugidou, C., Ouedraogo, I., 2014. First Report of Rice stripe necrosis virus Infecting Rice in Burkina Faso. Plant Disease 98, 1451–1451. 10.1094/PDIS-06-14-0626-PDN

Shepherd, D.N., Martin, D.P., Van Der Walt, E., Dent, K., Varsani, A., Rybicki, E.P., 2010. Maize streak virus: an old and complex ‘emerging’ pathogen. Molecular Plant Pathology 11, 1–12. 10.1111/j.1364-3703.2009.00568.x

Soullier, G., Demont, M., Arouna, A., Lançon, F., Mendez del Villar, P., 2020. The state of rice value chain upgrading in West Africa. Global Food Security 25, 100365. 10.1016/j.gfs.2020.100365

Tembo, M., Adediji, A.O., Bouvaine, S., Chikoti, P.C., Seal, S.E., Silva, G., 2020. A quick and sensitive diagnostic tool for detection of Maize streak virus. Sci Rep 10, 19633. 10.1038/s41598-020-76612-2

Tollenaere, C., Lacombe, S., Wonni, I., Barro, M., Ndougonna, C., Gnacko, F., Sérémé, D., Jacobs, J.M., Hebrard, E., Cunnac, S., Brugidou, C., 2017. Virus-Bacteria Rice Co-Infection in Africa: Field Estimation, Reciprocal Effects, Molecular Mechanisms, and Evolutionary Implications. Front. Plant Sci. 8, 645. 10.3389/fpls.2017.00645

Tra Bi, S.C., Dje, K.T., Coulibaly, T., Soumahoro, B., Tano, Y., 2020. Entomofaune du riz (Oryza sativa L.) en fonction des stades phénologiques dans un bas-fond, Daloa, Côte d’Ivoire. Afrique SCIENCE 16, 98–113.

Urbino, C., Thébaud, G., Granier, M., Blanc, S., Peterschmitt, M., 2008. A novel cloning strategy for isolating, genotyping and phenotyping genetic variants of geminiviruses. Virol J 5, 135. 10.1186/1743-422X-5-135

Varsani, A., Shepherd, D.N., Monjane, A.L., Owor, B.E., Erdmann, J.B., Rybicki, E.P., Peterschmitt, M., Briddon, R.W., Markham, P.G., Oluwafemi, S., Windram, O.P., Lefeuvre, P., Lett, J.-M., Martin, D.P., 2008. Recombination, decreased host specificity and increased mobility may have driven the emergence of maize streak virus as an agricultural pathogen. Journal of General Virology 89, 2063–2074. 10.1099/vir.0.2008/003590-0

Wang, P., Liu, J., Lyu, Y., Huang, Z., Zhang, X., Sun, B., Li, P., Jing, X., Li, H., Zhang, C., 2022. A Review of Vector-Borne Rice Viruses. Viruses 14, 2258. 10.3390/v14102258

